# Targeting pathogenic Lafora bodies in Lafora disease using an antibody-enzyme fusion

**DOI:** 10.1101/679407

**Authors:** M. Kathryn Brewer, Annette Uittenbogaard, Grant Austin, John J. McCarthy, Dyann M. Segvich, Anna DePaoli-Roach, Peter J. Roach, Bradley L. Hodges, Jill Zeller, James R. Pauly, Tracy McKnight, Dustin Armstrong, Matthew S. Gentry

## Abstract

Lafora disease (LD) is a fatal childhood epilepsy and a non-classical glycogen storage disorder with no effective therapy or cure. LD is caused by recessive mutations in the *EPM2A* or *EPM2B* genes that encode the glycogen phosphatase laforin and an E3 ubiquitin ligase malin, respectively. A hallmark of LD is the intracellular accumulation of abnormal and insoluble α-linked polysaccharide deposits known as Lafora bodies (LBs) in several tissues, including most regions of the brain. In mouse models of LD, genetic reduction of glycogen synthesis eliminates LB formation and rescues the neurological phenotype. Since multiple groups have confirmed that neurodegeneration and epilepsy result from LB accumulation, a major focus in the field has shifted toward the development of therapies that reduce glycogen synthesis or target LBs for degradation with the goal of treating LD. Herein, we identify the optimal enzymes for degrading LBs, and we develop a novel therapeutic agent by fusing human pancreatic α-amylase to a cell-penetrating antibody fragment. This antibody-enzyme fusion (VAL-0417) degrades LBs *in vitro*, shows robust cellular uptake, and significantly reduces the LB load *in vivo* in *Epm2a*-/- mice. VAL-0417 is a promising therapeutic for the treatment of LD and a putative precision therapy for an intractable epilepsy. Antibody-enzyme fusions represent a new class of antibody-based drugs that could be utilized to treat glycogen storage disorders and other diseases.

**One Sentence Summary:** An antibody-enzyme fusion delivering an amylase degrades the toxic polyglucosan bodies that cause Lafora disease, a fatal childhood epilepsy.

## Introduction

The progressive myoclonic epilepsies (PMEs) are a group of inherited disorders characterized by recurrent seizures, myoclonus, and progressive neurological decline. There are currently no treatments for PMEs, and anti-epilepsy drugs are palliative at best (*1*). Lafora disease (LD; epilepsy, progressive myoclonus type 2, EPM2) is a severe form of PME that typically manifests with tonic-clonic seizures and myoclonic jerks in the early teen years followed by rapid neurological deterioration, increasingly severe and frequent epileptic episodes, dementia and death within ten years of onset (OMIM: 254780). LD is caused by mutations in the *EPM2A* or *EPM2B* genes that encode laforin, a glycogen phosphatase, and malin, an E3 ubiquitin ligase that ubiquitinates enzymes involved in glycogen metabolism (reviewed in (*2*)). LD is distinguishable from other PMEs by the presence of cytosolic polysaccharide inclusions known as Lafora bodies (LBs) most notably in the brain, where they are found in neuronal cell bodies and dendrites and astrocytic processes, and in other tissues such as muscle, heart, and liver (*3−5*). Among the PMEs LD is uniquely considered a non-classical glycogen storage disease (GSD) (*6, 7*). Independent studies from multiple groups demonstrate that *Epm2a*-/- and *Epm2b*-/- mice recapitulate disease symptoms with respect to LB formation, neurodegeneration, neurological impairments, and susceptibility to epilepsy (*8−14*).

Glycogen and starch are the major carbohydrate storage molecules in mammals and plants, respectively (*15*). Mammalian glycogen, LBs, and starch are all glucose polymers (i.e. polysaccharides) composed of α-1,4-linked linear chains of glucose and α-1,6 linked branch points, but they have different properties. Glycogen consists of linear chains of ~13 glucose units that each give rise to two branches, producing a highly water-soluble molecule with a maximum diameter of 42 nm (*16*). LBs, which form in the absence of functional laforin or malin, are comprised of abnormal polysaccharide referred to as polyglucosan. The term polyglucosan refers to any glucose polymer that is abnormal, and other types of polyglucosan bodies have been described in some GSDs (*17*). Unlike glycogen, LBs are water-insoluble, hyperphosphorylated, contain long glucose chains, and range in size from a few microns to up to 35 or 40 μm (*18−21*). Dr. Gonzalo Rodriguez Lafora first described these inclusions in patient autopsies, noting their histochemical similarity to plant starch (*22*). Decades later Sakai and colleagues isolated these inclusions from patient tissue and showed that LBs are indeed structurally more similar to starch than to glycogen (*18, 19*). Plant starch is composed of infrequently branched, longer glucose chains that exclude water and allow for much larger, more energy-dense molecules to form (up to 100 μm in diameter) (*15, 23, 24*). Plant starch and LBs both contain significantly more covalently bound phosphate and longer glucan chains than mammalian glycogen (*19, 25, 26*). Loss of laforin or malin function results in the transformation of glycogen molecules into hyperphosphorylated polyglucosan molecules with an altered chain length distribution, and these aberrant molecules, prone to precipitation, accumulate to form LBs (*27*). How laforin and malin prevent this process is still being defined.

Results from multiple labs have established that accumulation of polyglucosan drives neurodegeneration in both LD and non-LD models. Glycogen synthase is the only mammalian enzyme able to catalyze glucose polymerization *in vivo. Epm2b*-/- mice lacking glycogen synthase (*Gys1)* in the brain are devoid of cerebral glycogen and LBs, exhibit no neurodegeneration, have normal hippocampal electrophysiology, and do not display an increased susceptibility to kainate-induced epilepsy (*28*). A similar phenotypic rescue was observed when *Gys1*-/- mice were crossed with *Epm2a*-/- mice (*29*). Furthermore, *Epm2b*-/- mice lacking just one *Gys1* allele in the brain have reduced glycogen and also show near complete rescue of these phenotypes (*28*). *Epm2a*-/- and *Epm2b*-/- mice lacking Protein Targeting to Glycogen (PTG), a protein that promotes glycogen synthesis, also exhibit reduced LB accumulation, and neurodegeneration and myoclonic epilepsy are resolved in these animals (*29−31*). These results demonstrate that decreased or complete absence of the glycogen synthesis machinery ablates LB formation, neurodegeneration, and epilepsy in LD mouse models. The reverse has also been observed: overexpression of a constitutively active form of glycogen synthase in otherwise wild type animals drives neurodegeneration in both flies and mice (*32*). The accumulating polysaccharide in transgenic animals overexpressing glycogen synthase is a polyglucosan (i.e. abnormal) rather than normal glycogen (*33*). Collectively, the aforementioned studies illustrate that cerebral LB accumulation is pathogenic. These studies have both elucidated the molecular etiology of LD and have made LBs an obvious therapeutic target (*34*).

These studies have motivated ongoing efforts to develop a targeted therapy (i.e. precision medicine) for LD (*35*). One form of precision medicine that has been successful with GSDs is the introduction of exogenous replacement enzymes. Enzyme replacement therapy has proven effective for Pompe disease (OMIM: 232300), an inherited GSD (*36*). Pompe patients are deficient in the lysosomal enzyme that degrades glycogen, acid α-glucosidase (GAA), and are currently treated with a recombinant human form of this protein known as rhGAA or alglucosidase alfa (Myozyme®, Lumizyme®, Genzyme) (*36, 37*). The uptake of rhGAA is a receptor-mediated endocytic process that targets rhGAA to the lysosome. However, there is a significant portion of cytosolic glycogen in Pompe patients, and since rhGAA only targets lysosomal glycogen, those with large pools of cytoplasmic glycogen do not respond well to the current therapy (*38*). In LD, LBs are entirely cytosolic. Histological studies from patient tissues report that LBs are not membrane bound, and this observation has been confirmed in mouse models (*11, 39−41*). Thus, a therapeutic enzyme degrading LBs must be delivered to the cytosol.

Although cytosolic targets remain challenging for protein therapeutics, antibody-based delivery platforms provide a means for penetrating the cell membrane (reviewed in (*42*)). The monoclonal anti-DNA autoantibody 3E10 and its antigen-binding (Fab) and variable domain (Fv) fragments can be fused to an enzyme to facilitate cytosolic delivery in multiple cell types (*43−45*). This antibody-enzyme fusion (AEF) strategy has proven successful for the replacement of a cytosolic enzyme and rescue of X-linked myotubular myopathy in mice (*46*). A Fab-GAA fusion has similar efficacy to reduce glycogen in Pompe mice with the additional advantage of targeting both lysosomal and cytosolic glycogen and is now in Phase 1/2 clinical trials (*47, 48*). Rather than utilizing the AEF platform to deliver a replacement enzyme, we generated an AEF designed to penetrate cells and degrade pathogenic LBs. Herein, we show that an AEF comprised of the humanized 3E10 Fab fragment and pancreatic α-amylase degrades LBs *in vitro*, degrades glycogen and polyglucosan *in situ*, and reduces LB load *in vivo*.

## Results

### Construction of an antibody-enzyme fusion (AEF) that degrades starch

An important consideration in designing an LB-degrading enzyme is that LBs are less sensitive to enzymatic hydrolysis than normal glycogen. Resistance to digestion with diastase (a generic term for enzymes that degrade polysaccharides) is a defining feature of LBs and one factor that differentiates LBs from glycogen (*40, 49*). However, in routine histological preparations, embedded sections are only incubated with diastase for 15-30 minutes prior to staining. Early studies reported that LBs in tissue sections can be digested by a 5-10 hour incubation with pancreatic α-amylase and γ-amylase (i.e. amyloglucosidase), but they are somewhat resistant to digestion by β-amylase or glycogen phosphorylase (*18, 19, 50*).

Glycogen, starch, and LBs are all polysaccharides with different properties. LBs are considered more similar to starch than glycogen due to their insolubility, elevated phosphate levels, and the increased chain length of their constituent glucose chains. We hypothesized that starch could serve as a proxy for LBs in a screen to identify a candidate therapeutic enzyme. We screened a panel of amylases to determine which possessed robust activity against starch, and therefore would likely degrade LBs. We found that only α-amylases showed significant activity, and pancreatic α-amylase had the highest capacity to degrade starch (Fig. 1A). Pancreatic α- amylase is naturally secreted by the human pancreas for digestion of carbohydrates in the gut, cleaving α-1,4 linkages to yield maltose, maltotriose and low molecular weight oligosaccharides. It has previously been shown to digest LBs in tissue sections (*50*). Thus, pancreatic α-amylase is a putative candidate for degrading LBs.

**Fig. 1.**
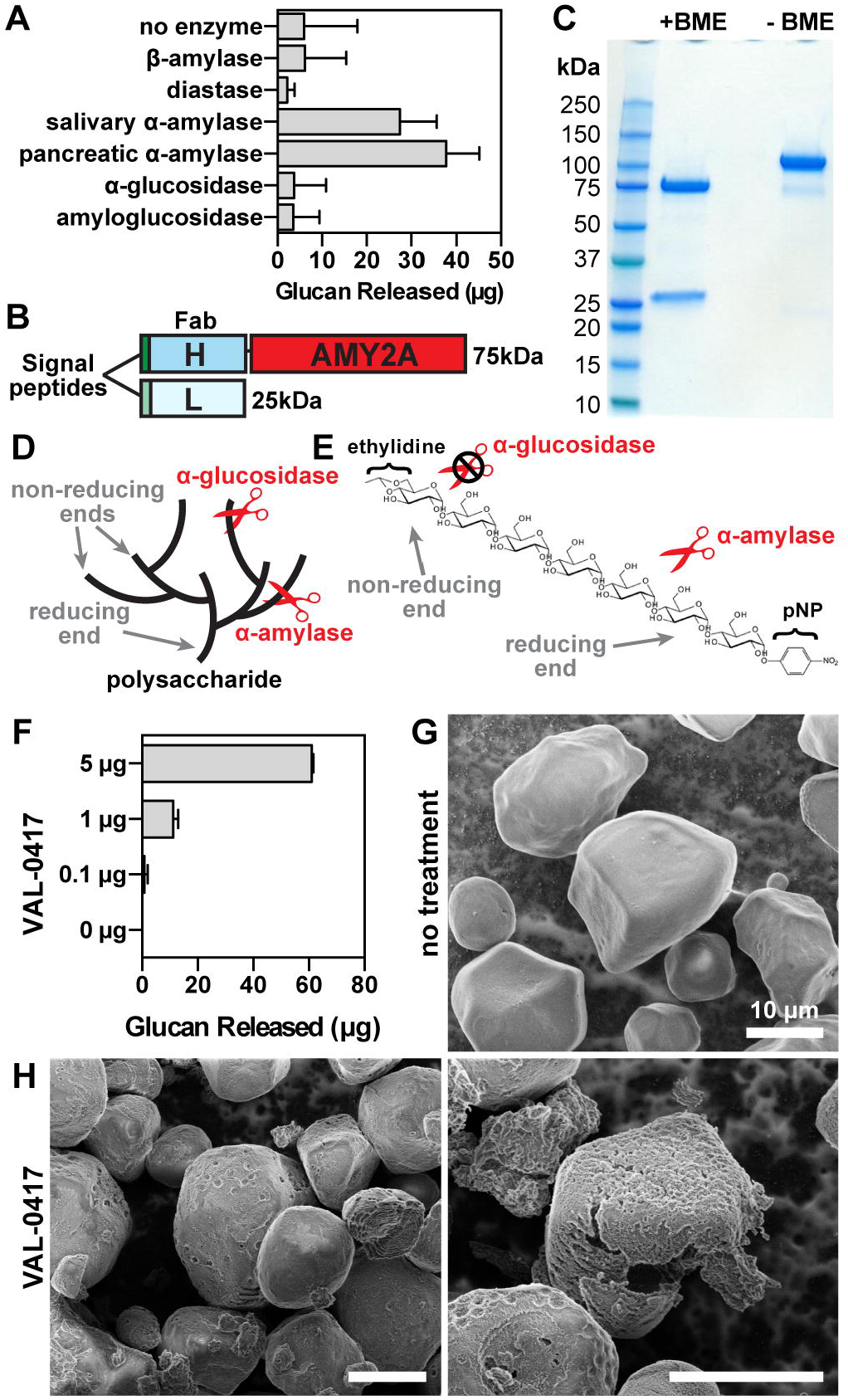
The antibody-enzyme fusion VAL-0417 degrades starch, a proxy for LBs. (A) Degradation of starch by a panel of amylases. 1 mg of starch was incubated with 5 μg of amylase overnight with constant agitation, then soluble and insoluble fractions were separated by centrifugation. Degradation product in the soluble fractions was quantified by measuring glucose equivalents (i.e. glucan). Reactions were performed in triplicate and data shown are means ± SD. (B) Schematic representation of VAL-0417. The gene encoding the human IgG1 Fab heavy chain fragment (H) was fused to AMY2A, encoding human pancreatic α-amylase, and coexpressed with the gene encoding the human light chain (L) in HEK293-6E cells. Heavy and light chain signal peptides were included to facilitate proper folding and assembly of the Fab fragment. The predicted molecular weight of each polypeptide is shown. (C) Purity of VAL-0417 assessed by reducing (+BME) and nonreducing (-BME) SDS-PAGE. (D) Illustration of glucan accessibility by α-glucosidase vs. α-amylase. Reducing (inner) and non-reducing (outer) ends of the polysaccharide are shown. (E) Chemical structure of the α-amylase-specific substrate E-G7-pNP. (F) Degradation of 1 mg starch after a 2-hour incubation with increasing amounts of VAL-0417, expressed as glucan released in the soluble fraction. Mean ±SD of triplicate reactions are shown. (G-H) Scanning electron micrographs of starch granules in the absence of treatment (G), and after overnight treatment with VAL-0417 (H). Samples were visualized at 2kV by an FE Quanta 250 scanning electron microscope.

An AEF designated VAL-0417 was generated by fusing the humanized 3E10 IgG1 Fab heavy chain fragment with human pancreatic α-amylase and then coexpressed in HEK293-6E cells with the corresponding light chain as a secreted heterodimer (Fig. 1B). The fusion was purified from HEK293-6E cell conditioned media by affinity chromatography, and the purity was determined by reducing and nonreducing SDS-PAGE (Fig. 1C). The addition of β- mercaptoethanol (BME) dissociated the heavy and light chains, producing two distinct bands of 75 and 25 kDa; in the absence of BME a single 100 kDa band was observed, corresponding to the intact Fab-amylase fusion. The specific activity of an α-amylase can be tested using a substrate that consists of seven glucose moieties and a chromophoric *pa*ra-nitrophenyl moiety that is capped with an ethylidine group to block the action of other hydrolytic enzymes (Fig. 1D and E) (*51*). Using this substrate, the specific activity of VAL-0417 was determined to be 10.5 units/mg.

VAL-0417 activity was then tested against plant starch. After a 2-hour incubation, VAL-0417 degraded starch in a dose-dependent manner: 5 μg of enzyme released 61 μg of soluble glucan product (Fig. 1F). We employed scanning electron microscopy (SEM) to visualize the effects of degradation on the starch granules. In the absence of enzymes, starch granules had a smooth, polyhedral or subspherical appearance (Fig. 1G). Overnight treatment with VAL-0417 produced cavities in the granule surface, representing areas of active hydrolysis, and numerous partially degraded granules were visible (Fig. 1H). When the ratio of VAL-4017 to starch was reduced by 10-fold, amylolysis was less extensive: fewer partially degraded particles were visible and most granules were only decorated with small pits (fig. S1). Other groups have similarly observed “pitting” of starch granules with α-amylase and other amylolytic enzymes (*52, 53*).

### Isolation and characterization of LBs from LD mice

Polysaccharides (both glycogen and LBs) are typically purified from *Epm2a*-/- and *Epm2b-/*- mice based on a method described by Pflüger in 1909 where the tissue is heated in strong KOH and the glycogen is precipitated with alcohol (*54, 55*). The isolated particles are 15-65 nm in size when visualized by transmission electron microscopy (*20*), orders of magnitude smaller than the intact LBs observed in fixed tissue sections, and the method does not separate LBs from glycogen. It is well-established that heating changes the physiochemical properties of starch, leads to granule swelling, and enhances its sensitivity to enzymatic degradation (*56*). Therefore, it is likely that this protocol disrupts the native structure of LBs, intermixing the individual polyglucosan constituents of LBs with normal glycogen molecules. We developed a procedure for isolating pure and native LBs from tissue so that we could test the sensitivity of LBs to VAL-0417 degradation.

A study from the 1960s showed that native LBs could be isolated from patient brain tissue by centrifugation steps and successive treatments with proteolytic enzymes (*18*). We designed a similar method for isolating native LBs from the brain, heart and skeletal muscle of *Epm2a*-/- mice (Fig. 2A, see Materials and Methods). Our procedure separates soluble glycogen (present in the supernatant) from the insoluble LBs (fig. S2 and Supplementary Methods). Final LB yields from each tissue type corresponded to ~30% of the total polysaccharide detected in the original tissue homogenate (Fig. 2B and C). Phosphate content of the final LBs from skeletal muscle was 5-fold higher than phosphate in normal muscle glycogen, which is in agreement with published results (Fig. 2D) (*20*). Iodine spectral scans of the purified LBs were compared to those of commercial glycogen and amylopectin, the major component of plant starch to which LBs are most similar. While glycogen has a peak maximum at ~440 nm when stained with Lugol’s iodine, amylopectin has a right-shifted spectrum and a peak maximum at 560 nm due to its reduced branching (*57*). The spectra of the purified LBs were also shifted toward longer wavelengths, with peak maxima at 510-520 nm (Fig. 2E). These data indicate that the purified LBs are less branched than glycogen, consistent with previous reports on *Epm2a*-/- and *Epm2b*-/- mice (*10, 20*).

**Fig. 2.**
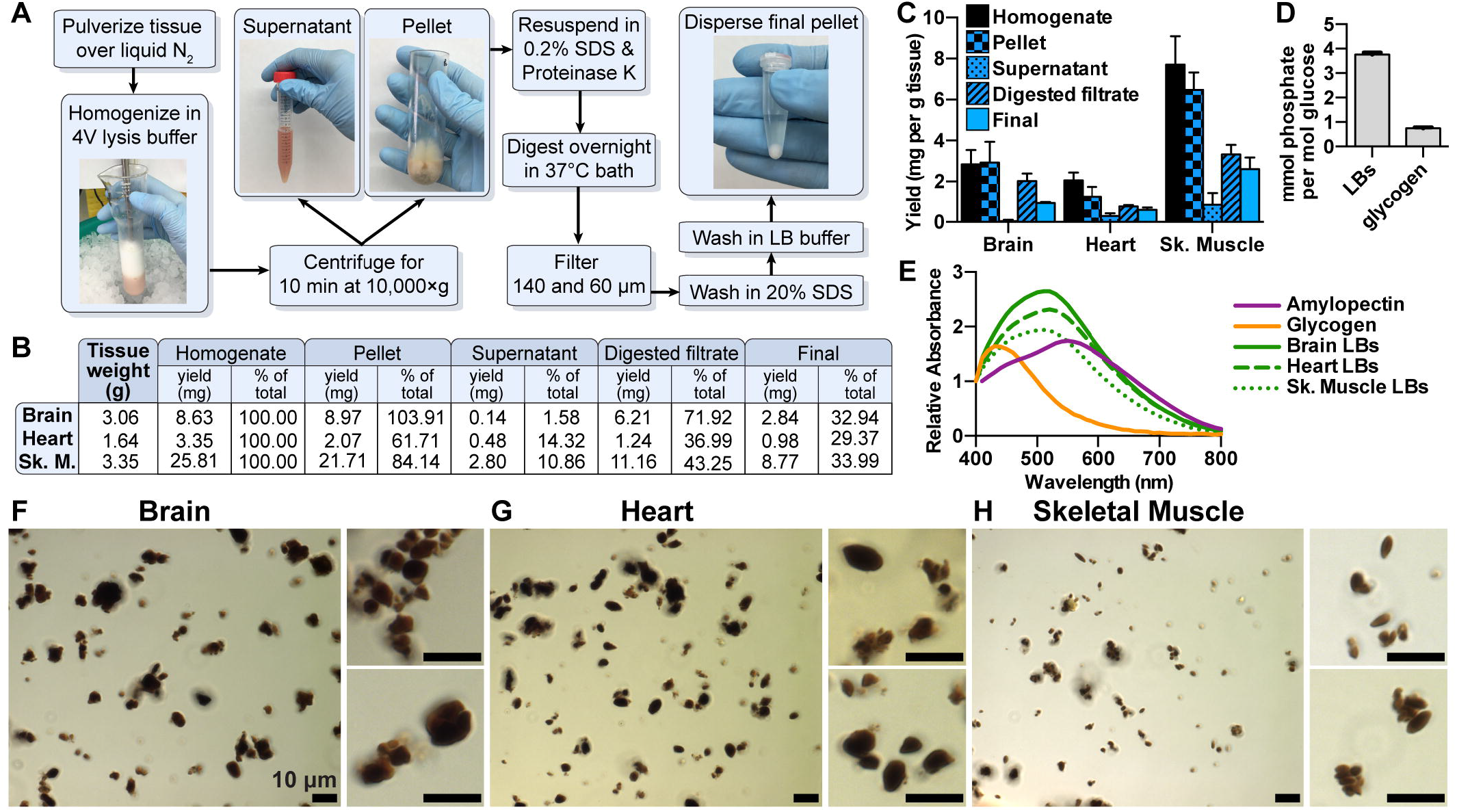
A novel protocol for isolating native LBs from LD mice. (A) LB purification scheme. (B) Polysaccharide was purified at different steps in the protocol via the Pflüger method and quantitated via glucose measurement following hydrolysis. Initial tissue weights and total polysaccharide at each step are shown. (C) Polysaccharide yields normalized to tissue weight. Triplicate samples were removed from each fraction and each measured in triplicate. Mean ±SD are shown. (D) Phosphate content of LBs from skeletal muscle and normal rabbit muscle glycogen. Mean ±SD of triplicate measurements are shown. (E) Normalized iodine spectra of purified LBs compared to commercial liver glycogen and amylopectin. Spectra shown are an average of 3 replicates. (F) Brain, (G) heart, and (H) skeletal muscle LBs stained with Lugol’s solution and visualized using a Zeiss Axioimager Z1.

Final LB preparations were also stained with Lugol’s iodine and examined by light microscopy. Minimal tissue debris was observed and the LBs tended to clump (Fig. 2, F to H). LBs from different tissues also had noticeably distinct morphologies. Brain LBs were generally spherical, sub-spherical or irregularly shaped (Fig. 2F), and a range of sizes were observed: the maximum diameter of a single LB ranged from 1-10 μm, most being 2-4 μm (fig. S3A). These preparations also had a significant amount of granular material that stained less intensely with iodine; small amounts of this material were visible throughout the slide, but appeared to form a large aggregate in some fields of view (fig. S3D). Similar PAS-positive granular material has been reported in fixed tissue sections from patient brain biopsies or LD mice, which are referred to as dust-like particles, or simply “dust” (*40, 58−60*).

Heart LBs were more often ellipsoid or ovoid in shape, frequently with pointed ends, although some were sub-spherical like the brain LBs (Fig. 2G). Heart LBs also ranged in diameter/length from 1-10 μm, most of them being 2-4 μm (fig. S3B). Dust-like particles were also observed among the heart LBs, although no masses of granular material were observed as with the brain LBs (fig. S3E. LBs have been reported in LD patient cardiac tissue: in fixed and stained tissue sections, they are irregularly shaped or elongated with pointed ends (*59, 61−63*). Although these reports do not discuss dust-like particles in the heart, they appear to be present in the micrographs, particularly in the studies that involve immunostaining with an LB-specific antibody (*61, 62*). PAS-stained cardiac sections from malin KO mice show LBs of similar shape and size and diastase-resistant granular material (*9, 11*). LBs from skeletal muscle were the smallest and most homogenous in size and shape: the vast majority were ellipsoid and 1-3 μm long (Fig. 2H, fig. S3C). There appeared to be virtually no dust in these preparations. In LD patients, LBs have been reported to similarly range in diameter from 0.2-2 μm in skeletal muscle (*58*).

We also utilized SEM to visualize the purified LBs, which could be captured at high magnification (Fig. 3, fig. S4). LBs from all three tissues had a similarly textured appearance, in contrast to the smooth surface of starch (Fig. 1G). As observed via light microscopy, individual brain LBs up to 10 μm in diameter were observed, which were spherical or irregularly shaped (Fig. 3A). Ovoid and ellipsoid LBs from heart and small ellipsoid LBs from skeletal muscle were also consistent with our light micrographic observations (Fig. 3, B and C).

**Fig. 3.**
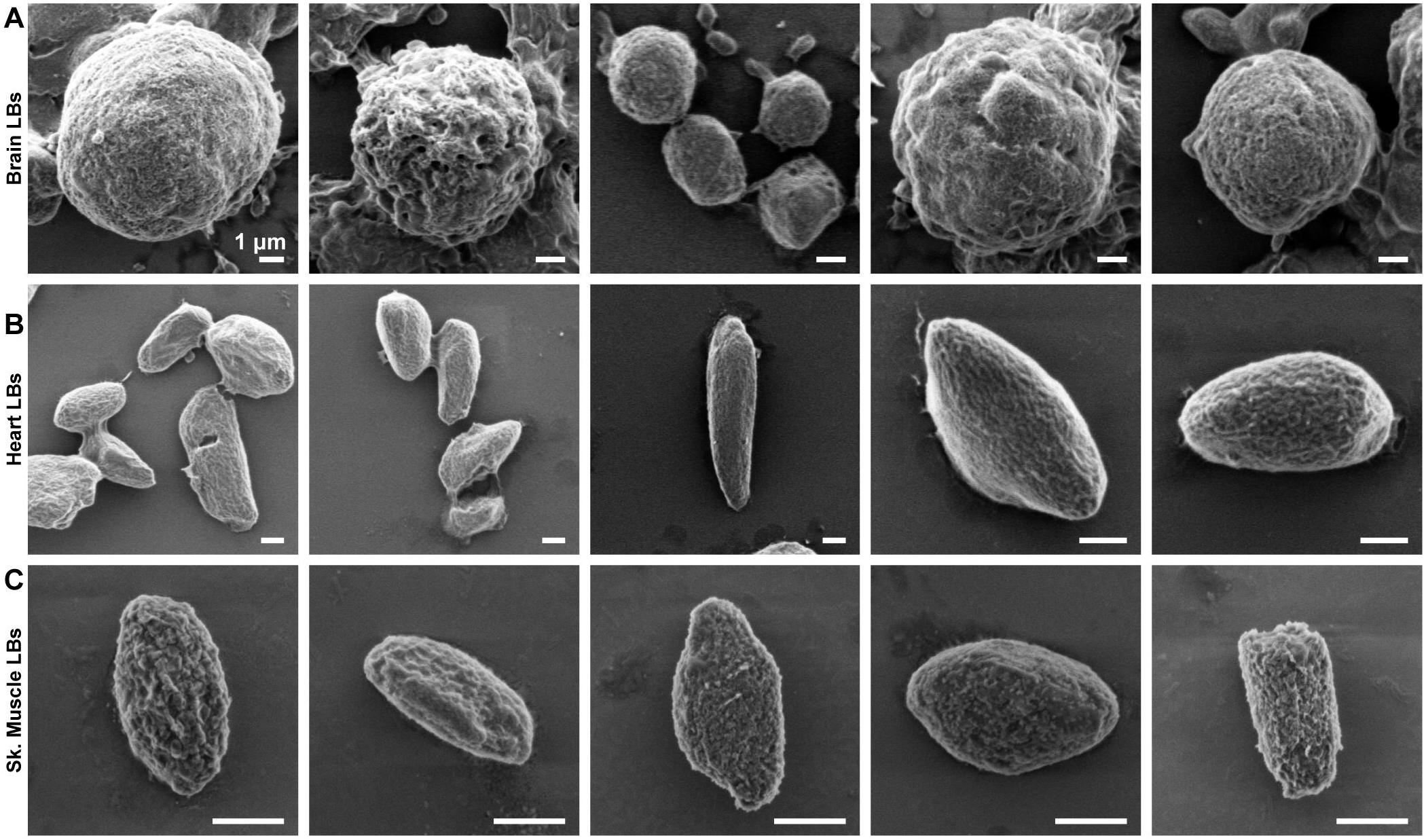
Scanning electron micrographs of isolated LBs. Scanning electron micrographs of LBs purified from brain (A), heart (B), and skeletal muscle (C). Samples were visualized at 2 kV using an FE Quanta 250.

### Degradation of LBs by VAL-0417

Native LBs purified from different tissues possess unique morphology. To determine if VAL-0417 could degrade the different types of LBs, we incubated 50 μg of *Epm2a*-/- LBs from brain, heart and skeletal muscle with VAL-0417 and observed a consistent dose-response effect: 1 μg VAL-0417 released 6-9 μg of glucan into the soluble fraction, indicating 12-18% of the LBs had been degraded (Fig. 4A). Increasing the dose to 10 μg of VAL-0417 released over twice as much, 15-19 μg of glucan, corresponding to 30-38% of the total substrate. We also purified LBs from skeletal muscle of 12-month-old *Epm2b*-/- mice and found they were indistinguishable from *Epm2a*-/- skeletal muscle LBs in appearance, size and iodine spectra (fig. S5). We observed a comparable dose-response effect of VAL-0417 on *Epm2b*-/- LBs: 1 μg and 10 μg VAL-0417 released 16% and 32% of the total, respectively (Fig. 4A). In a similar experiment, we incubated LBs with 10 μg VAL-0417 overnight and then visualized the product by staining the insoluble fraction with Lugol’s iodine. Untreated LBs had no change in appearance and seemed fully intact after the overnight agitation, but VAL-0417 treatment led to fewer iodine-positive LBs and the appearance of a filamentous degradation product (Fig. 4B).

**Fig. 4.**
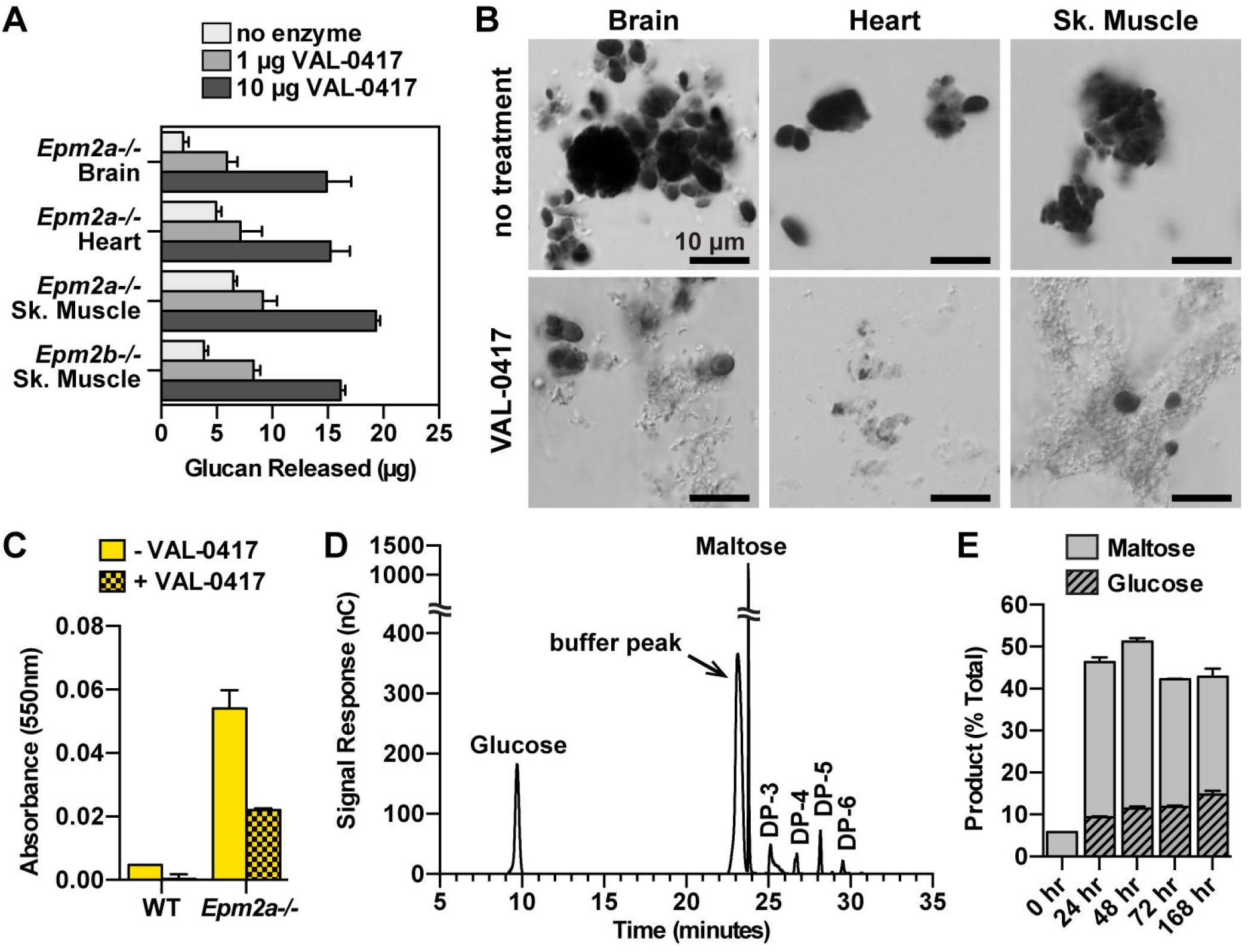
VAL-0417 degrades LBs *in vitro*. (A) Degradation of 50 μg LBs from different tissues after overnight incubation with 0, 1, or 10 μg VAL-0417. (B) LBs from different tissues after overnight incubation +/- 10 μg VAL-0417. Insoluble fractions were resuspended post-degradation, stained with Lugol’s solution, and visualized using a Nikon Eclipse E600. (C) Iodine absorbance in pellet fractions after incubation of WT and *Epm2a*-/- skeletal muscle homogenates +/- 25 μg VAL-0417. In (A), (C), and (E) mean ± SD of triplicates are shown. (D) HPAEC-PAD chromatogram of degradation product after incubation with VAL-0417 for 168 hours. Peak identities were determined based on the elution profile of degradation buffer and glucan standards. The chromatogram is representative of triplicate experiments, and glucose/maltose quantification from the replicates is shown in (E). (E) Quantification of glucose and maltose at various time points throughout the degradation reaction of LBs with VAL-0417. Product released are expressed as a percentage of total LBs.

Previous work has shown that various proteins are associated with LBs *in vivo (11, 60*). Our purification protocol removes these proteins, so it is possible that an intact protein coat could inhibit LB degradation by VAL-0417. To test this possibility, we measured LB load in crude muscle homogenates after incubation with VAL-0417 using iodine absorbance. While virtually no absorbance was detected at 550nm in the WT samples with or without VAL-0417 treatment, treatment of the *Epm2a*-/- samples with VAL-0417 reduced iodine absorbance by >50%, indicating substantial LB degradation (Fig. 4C).

Since VAL-0417 is a potential therapeutic, it is important to define the soluble product(s) it releases from LBs. Pancreatic α-amylase is reported to release glucose, maltose, maltotriose and other short oligosaccharides (*64*). Amylolytic products can be separated and quantified via high-performance anion-exchange chromatography coupled with pulsed amperometric detection (HPAEC-PAD). We digested LBs from *Epm2a*-/- skeletal muscle with VAL-0417 for a total of 168 hours with additional enzyme added every 24 hours. At 24, 48 and 72 hour time points, the soluble fractions of LB digestions were removed for HPAEC-PAD analysis. At all 4 time points, maltose and glucose were the major degradation products, followed by small amounts of oligosaccharides with 3, 4, 5 and 6 degrees of polymerization (DP-3, 4, etc., where each DP corresponds to one glucose unit) (Fig. 4D, fig. S7). Quantitation of the glucose and maltose peaks showed that maltose levels remained constant throughout degradation and glucose levels increased steadily over the 168-hour experiment (Fig. 4E). Cumulatively, these data demonstrate that VAL-0417 robustly digests brain, heart and muscle LBs *in vitro*. It is not inhibited by endogenous proteins in muscle homogenates, and its degradation products are primarily glucose and maltose, with a few short oligosaccharides.

### VAL-0417 uptake and polysaccharide degradation *in situ*

To degrade LBs *in vivo*, VAL-0417 must penetrate and remain active inside cells. Cell penetration of the parent 3E10 mAb and its fragments require functional expression of the membrane equilibrative nucleoside transporter 2 (ENT2), which is involved in a nucleoside salvage pathway and is ubiquitously expressed in rodents and humans (*65−67*). VAL-0417 uptake and activity were assessed in two cell lines: fibroblast-derived Rat1 cells accumulating normal glycogen and HEK293 cells engineered to accumulate polyglucosan (HEK293-PTG/PP1Cα) (fig. S6; see Supplementary Methods). After a 20-hour treatment with increasing doses of VAL-0417, cells were washed, lysed, and probed for ENT2 and VAL-0417. ENT2 protein levels were stable in both lines, though Rat1 cells have slightly higher levels, and were unchanged with increasing concentrations of VAL-0417 (Fig. 5A). Western analysis of VAL-0417 using an antibody for AMY2A revealed a dose-dependent increase in protein levels (Fig. 5A). Polysaccharide (glycogen in Rat1 cells and polyglucosan in HEK293-PTG/PP1Cα cells) in the treated cells was also measured. Both Rat1 and HEK293-PTG/PP1Cα lines displayed a dosedependent decrease in total polysaccharide, reductions of 30% and 26%, respectively (Fig. 5B). Thus, VAL-0417 is both taken up by cells and degrades glycogen and polyglucosan *in situ*.

**Fig. 5.**
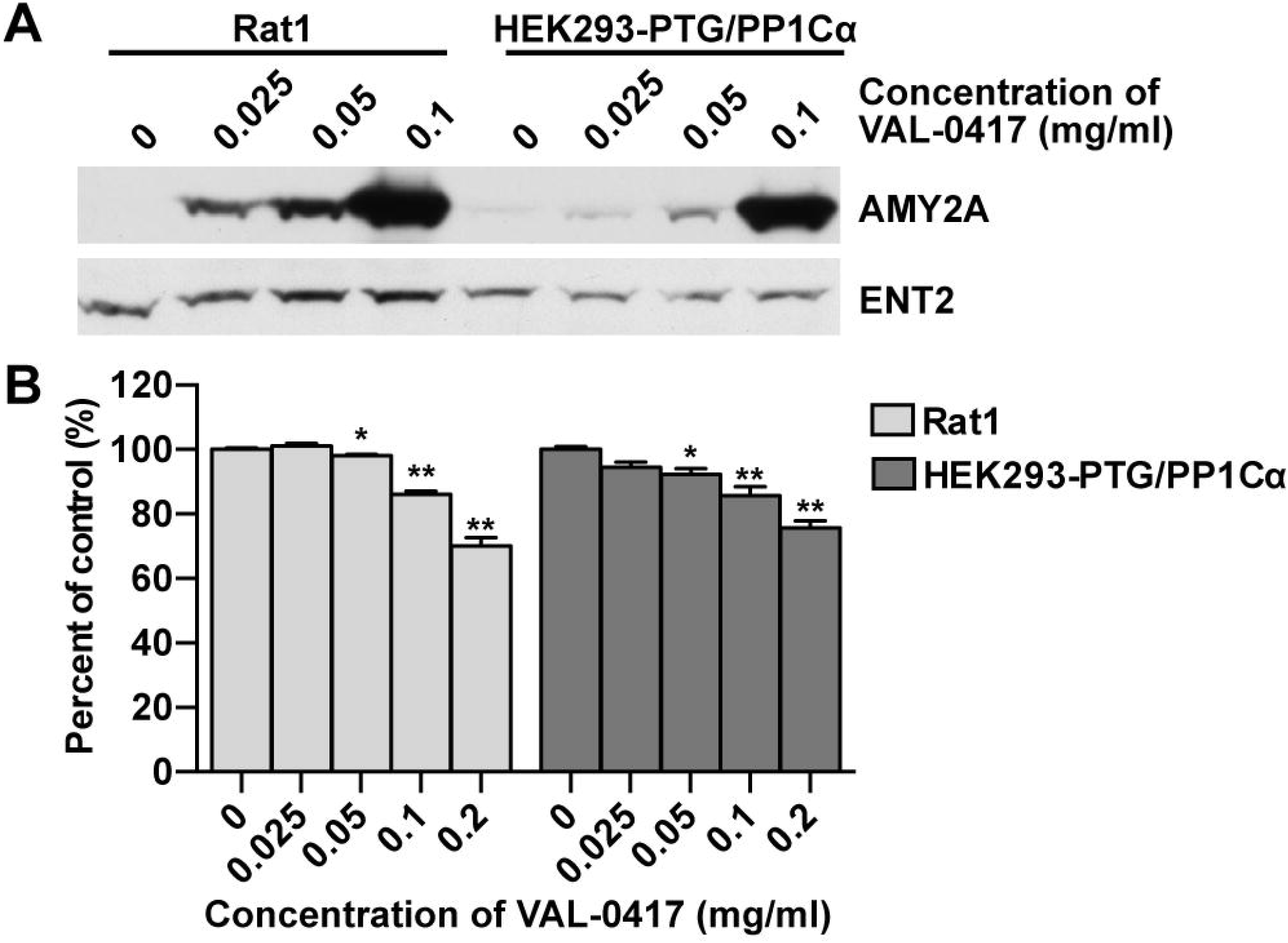
VAL-0417 uptake and polysaccharide reduction in cell culture. (A) Rat1 cells and HEK293 cells stably expressing PTG and PP1Cα after 20-hour treatment with increasing concentrations of VAL-0417. VAL-0417 levels were assessed by Western blotting using an anti-AMY2A antibody. ENT2 levels were also quantified by Western blotting. (B) Polysaccharide levels in Rat and HEK293-PTG/PP1Cα cells after 20-hour treatment with VAL-0417, expressed as a percent of controls. Data shown are a mean of 4 measurements ± SE. Statistical significance is indicated: \**p* ≤ 0.05, \*\**p* ≤ 0.01, *** *p* ≤ 0.001.

### *In vivo* uptake of VAL-0417 and LB reduction in *Epm2a*-/- mice

We have shown that VAL-0417 degrades LBs *in vitro*, penetrates cultured cells, and degrades glycogen and polyglucosan *in situ*. We next tested VAL-0417 uptake in an *in vivo* setting and biodistribution after peripheral injections. We performed intramuscular (IM) injections of VAL-0417 to determine if it can be detected in the injected muscle and other tissues and if it is quickly degraded and/or cleared. We injected 0.6 mg VAL-0417 into the left gastrocnemius muscles of WT C57BL/6 mice, then 2 and 24 hours post-injection, euthanized the mice and collected right and left gastrocnemii, right and left quadriceps, heart, liver, and brain. The same tissues were also collected 2 hours post-injection from control mice treated with phosphate-buffered saline (PBS). We designed a sandwich enzyme-linked immunosorbent assay (ELISA) to specifically detect VAL-0417 in tissue homogenates with very low background. At the 2-hour VAL-0417 post-injection time point, 3200 ng of VAL-0417 per mg protein (15% of the injected amount) was detected in the injected (left) gastrocnemius (Fig. 6A). After 24 hours, VAL-0417 levels dropped by 20-fold (to 151 ng per mg protein), 0.7% of the initial injected amount. At 2 hours, low levels of VAL-0417 (2-9 ng per mg protein, 0.02-0.1% of the injected amount) were detected in the other muscles, heart, liver, and brain, indicating some VAL-0417 had entered circulation. After 24 hours, VAL-0417 levels nearly returned to baseline in muscles, liver, and brain, but remained constant in the heart. Since blood circulates the entire body of a mouse within minutes, the very high levels of VAL-0417 in the injected muscle 2 and 24 hours postinjection strongly suggest that VAL-0417 is taken up by cells rather than being immediately cleared. The low levels of VAL-0417 in non-injected tissues indicates VAL-0417 entered the circulation. In mice, ENT2 expression levels are high in brain, muscle and heart, and low in liver (*66*). It is possible that these tissues took up a small amount of VAL-0417 from the circulation that was mostly cleared by 24 hours. ENT2 levels were probably not limiting since VAL-0417 levels were similar regardless of tissue differences in ENT2 expression. It is notable that VAL-0417 levels remained constant in heart even after 24 hours.

**Fig. 6.**
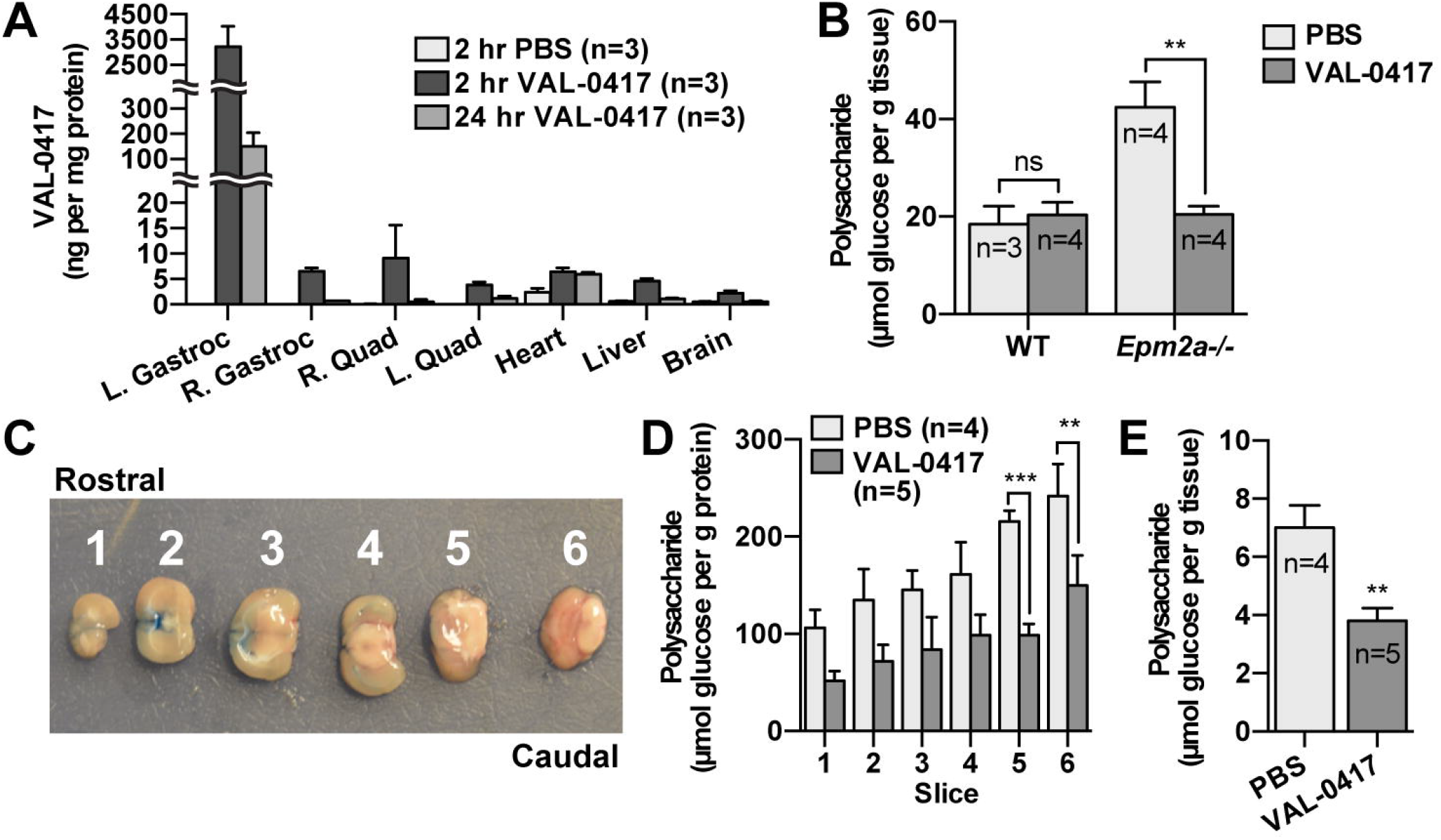
VAL-0417 reduces LB load *in vivo*. (A) Biodistribution of VAL-0417 levels determined by sandwich ELISA after IM injection of WT mice with PBS or 0.6 mg VAL-0417. (B) WT and *Epm2a*-/- mice were given 3 IM injections of PBS or VAL-0417 over the course of one week. Polysaccharide in the injected gastrocnemii was isolated via the Pflüger method and quantified by glucose measurement following hydrolysis. (C) Representative photograph of six 2.0 mm coronal sections. Blue dye was injected through the cannula into the lateral ventricle to verify its location within slice 2. (D) Polysaccharide content normalized to protein content in *Epm2a*-/- brain slices after 28 days of continuous ICV infusion of VAL-0417 or PBS. (E) Total brain polysaccharide normalized to tissue weight of PBS and VAL-0417 treated *Epm2a*-/- mice. In (B), (D), and (E), at least three technical replicates were performed for each isolated tissue/slice to determine an average per tissue/slice per animal, and data shown are the mean from each treatment group ± SE. The numbers of animals in each treatment group (n) are shown. Statistical significance is indicated: ns *p* > 0.05, \**p* ≤ 0.05, \*\**p* ≤ 0.01, *** *p* ≤ 0.001.

We next tested whether IM injections of VAL-0417 reduced polysaccharide levels *in vivo* in *Epm2a*-/- mice. To determine polysaccharide levels in individual muscles, we utilized the Pflüger method rather than our native LB protocol because the Pflüger method recovers ~100% of the polysaccharides, enables more accurate and sensitive measurements from small amounts of tissue, and yields both normal glycogen and LBs. 10-month-old WT and *Epm2a*-/- mice were injected in the right gastrocnemius with either 0.6 mg VAL-0417 or PBS. Three injections were performed over the course of one week, on days 1, 4, and 7, and on day 8 the mice were euthanized and the injected muscles were collected for total polysaccharide quantification. Polysaccharide levels in the WT mice were similar regardless of treatment, and consistent with reported skeletal muscle glycogen levels for C57BL/6 mice (*20*). In *Epm2a*-/- animals, VAL-0417 treatment reduced polysaccharide levels by 53% relative to the PBS-treated controls (Fig. 6B). These data indicate that *in vivo*, VAL-0417 does not decrease normal glycogen levels, but degrades LBs in *Epm2a*-/- mice. Since polysaccharide levels in the *Epm2a*-/- mice include LBs, and LBs are cytosolic, these data indicate that VAL-0417 penetrates cells, remains active, and degrades LBs *in vivo*.

Cerebral LB accumulation drives epilepsy and neurodegeneration in LD (*28*). Intracerebroventricular (ICV) administration enables drug delivery directly to the ventricles of the brain, bypassing the blood-brain barrier. It is considered a safe and well-tolerated route of drug administration in both pediatric and adult populations (*68*). To target cerebral LBs, we administered VAL-0417 directly into the cerebral lateral ventricle of 5-month-old *Epm2a*-/- mice using a surgically implanted ICV catheter attached to an osmotic pump. A continuous infusion of VAL-0417 (0.08 mg/day) or PBS was administered for 28 days via the ICV pump. Mice were then euthanized and brains were removed and sectioned as shown (Fig. 6C). Polysaccharides were isolated from each section, quantified, and normalized to protein levels. Polysaccharide levels relative to protein content were lowest in the rostral section and highest in the caudal section (Fig. 6D). This is the first time polysaccharide distribution across different brain regions has been quantitated in *Epm2a*-/- mice, and is consistent with the high levels of LBs observed in histological sections from brainstem and cerebellum (*3, 10, 11*). Treatment with VAL-0417 reduced the polysaccharide load in all sections, with statistical significance in the two most caudal sections that contain the highest glucan load (Fig 6D). We also quantified whole brain polysaccharide relative to tissue weight (Fig. 6E). Polysaccharide levels in PBS-treated animals are consistent with what has been previously reported for *Epm2a*-/- animals of this age, i.e. 3-4 times higher than WT levels (2 μmol glucose/g tissue) (*20, 69*). Treatment with VAL-0417 reduced total brain polysaccharide in the *Epm2a*-/- mice by 43% (Fig. 6E), only 2-fold higher than normal brain glycogen levels.

### Discussion

LD is caused by mutations in one of two genes both encoding enzymes that impact glycogen structure. Efforts from multiple labs have shown that LBs are the pathogenic epicenter of LD and removal of LBs has become a therapeutic target for treating LD (*34*). In this study, we combine insights from the fields of antibody-based therapeutics, GSDs, carbohydrate metabolism, and LD to design an AEF that degrades LBs *in vitro*, penetrates cells and reduces polysaccharide levels *in situ*, and decreases polysaccharide load *in vivo* via both IM and ICV injection. This is the first time a significant LB reduction has been achieved in adult LD mice by the use of a drug.

Antibody-based drugs have made a major impact on the treatment of cancer, autoimmune and inflammatory diseases with worldwide revenues of ~$89 billion in 2016 (*70*). Antibody-based drugs are typically whole IgGs designed to bind and block receptors, deplete target cells, downregulate a receptor or initiate signaling. Bispecific antibodies, antibody fragments and antibody-drug conjugates (ADCs) are also being developed across multiple diseases, further expanding the diversity of therapeutics utilizing antibody targeting (*70−73*). Herein, we report a novel class of antibody-based drugs with a similar construction to ADCs: antibody-enzyme fusions (AEFs). Rather than delivering a cytotoxin, AEFs deliver an enzyme to a specific target. The antibody or antibody fragment is essential both for targeting and cellular penetration. AEFs have recently been reported for the treatment of two congenital disorders: a muscle wasting disorder known as X-linked myotubular myopathy and the GSD Pompe disease. In the first study, the deficient lipid phosphatase myotubularin fused to an Fv fragment was delivered by intramuscular injection and improved muscle function in a mouse model of myotubular myopathy (*46*). In the second, a Fab fragment fused to GAA delivered intravenously reduced glycogen load and alleviated pathology in GAA-deficient mice (*47*). These two studies illustrate that enzyme replacement therapy, which can be problematic for cytosolic enzymes, can be enhanced by fusion of the replacement enzyme with a cell-penetrating antibody fragment. We now demonstrate that VAL-0417, an AEF comprised of a Fab fragment fused to pancreatic α- amylase, degrades LBs and reduces polysaccharide loads *in vivo*. Unlike the previous two studies, VAL-0417 does not replace a missing enzyme, but rather delivers a natural human enzyme to a new target. Since LBs are structurally similar to plant starch, and α-amylase naturally degrades dietary starch in the human digestive tract, α-amylase is a logical choice for the VAL-0417 payload.

It is important to distinguish normal glycogen from polyglucosan. Polyglucosan bodies (PGBs) have historically been defined as “small, non-membrane-bound cytoplasmic structures composed largely of unusual glucose polymers” (*33*). They are typically a pathological feature of GSDs, although PGBs known as corpora amylacea are a hallmark of normal aging and neurodegenerative disorders (*17*). PGBs are not found in all GSDs, and not all PGBs are identical with respect to physiochemical structure. The abnormal polyglucosan found in Cori’s disease is characterized by short outer chains, while the polyglucosan of LD, Anderson disease, Adult Polyglucosan Body Disease, and Tarui disease contain very long chains (*33, 74*). The Pflüger method, originally designed to purify normal glycogen, also precipitates polyglucosan. Herein we refer to Pflüger-isolated material as polysaccharide, since it contains both glycogen and polyglucosan. Furthermore, the Pflüger method includes a boiling step that solubilizes polyglucosan and destroys LB superstructure, which is not included in our LB purification procedure. LBs (and PGBs) may be considered a higher-order structure of polyglucosan. The LB purification protocol we designed does not destroy the native LB structure and separates LBs from normal, soluble glycogen.

LBs from the different tissues displayed varied morphologies and subtle differences in iodine spectra. Brain LBs were the most irregular in size and shape, included dust-like particles, and reached the largest size. LBs in the heart were also surprisingly large and included some dust-like particles. LBs from skeletal muscle appeared well organized: they were very small and homogenous in shape, and no dust-like particles were present. Although LBs from all 3 tissues had iodine spectral peak maxima at similar wavelengths, indicative of similar degrees of branching, the higher amplitude of the spectra of brain and heart LBs suggests that they contain longer chains compared to LBs from skeletal muscle (*57, 75*). Indeed, Nitschke *et al*. showed using HPAEC-PAD that in *Epm2a*-/- and *Epm2b*-/- mice, brain glycogen had a more exacerbated shift in chain length distribution than muscle glycogen (*69*). Although the brain is most severely affected in LD and a neurological phenotype predominates, heart failure and arrhythmia have been reported in LD patients, and *Epm2a*-/- and *Epm2b*-/- mice have metabolic cardiomyopathy (*76−78*). Glycogen metabolism is variable with tissue type: glycogen synthase and glycogen phosphorylase have tissue-specific isoforms that are disparately regulated by cellular signals (*79*); the ultrastructure of glycogen is different under different metabolic conditions and in different organs (*80, 81*); subcellular glycogen pools are variable (*16*); and both phosphate levels and average chain length differ with species and tissue type (*82, 83*). Due to these differences, the LBs may be more pathological in brain and heart than in skeletal muscle. Conversely, different cell types may be more sensitive to the presence of the LBs, and/or the metabolic perturbations induced by the LBs may have distinct consequences in different organs. Tissuespecific differences are important to consider throughout the clinical development of VAL-0417. ICV administration of VAL-0417 may attenuate neurological symptoms, but cardiac, liver and/or muscular pathology may require treatment by intravenous or intramuscular injection of VAL-0417.

We found that VAL-0417 primarily releases maltose and glucose from LBs, and glucose levels gradually increased over time, as expected for an α-amylase. Cytosolic free glucose would likely be phosphorylated to produce glucose-6-phosphate, entering central carbon metabolism. Although maltose is not considered a typical participant in intracellular energy utilization, studies have shown that maltose can be transported into and out of cells and metabolized in cell culture and humans after intravenous infusion (*84, 85*). The small oligosaccharides (DP-3, DP-4, etc.) would likely remain cytosolic and inert until they are further metabolized to glucose and maltose. Importantly, IM injections of VAL-0417 in *Epm2a*-/- mice reduced the polysaccharide load (i.e. LBs and glycogen) to WT levels, but did not decrease glycogen levels in WT mice. In contrast, we had observed that VAL-0417 reduced glycogen levels in cell culture. It should be noted that the cell culture studies were performed in high glucose media, known to promote glycogen accumulation in cultured cells. Thus, it is possible that a reduction in polysaccharides with VAL-0417 treatment is only detected when the polysaccharide accumulates above a baseline level, as glycogen accumulates in cultured cells or LB in LD tissues. When glycogen levels drop below a certain threshold due to degradation by VAL-0417, the cellular machinery may respond to restore those levels. Glycogen is likely to be maintained at a certain baseline level *in vivo* in muscle, and this may explain why no change was observed in WT muscle treated with VAL-0417. The observation that VAL-0417 reduced polysaccharide levels in *Epm2a*-/- muscle and brain indicates LBs are degraded *in vivo*. We have initiated further studies to define the metabolic changes invoked by VAL-0417 in WT and LD mice.

Given these exciting results, we are now performing pre-clinical experiments with VAL-0417 to define the minimal *in vivo* efficacious dose, the therapeutic window for treatment, and the necessary dosing frequency. We have established a robust VAL-0417 ELISA-based quantitation method that will be utilized to define the pharmacokinetic and pharmacodynamic parameters. Further, we are actively developing biomarkers that could be used to assess VAL-0417 treatment efficacy and monitor ongoing therapy. In the current study, we observed dramatic effects with VAL-0417 in brain of 6-month old mice and muscle of 10-month old mice. Moving forward, we will define the range of doses that most effectively clear LBs in animals at various ages as well the dose that effectively prevents the appearance of LBs in younger animals.

We have demonstrated that VAL-0417 is a putative precision therapy for LD, an intractable epilepsy. Currently, >30% of epilepsies, including LD, are resistant to available antiseizure drugs (ASDs) and are known as refractory epilepsies (*86*). More than 35 different iterations of ASDs have been discovered over the past 150 years with most of them acting primarily on ion channels or neurotransmitters. Due to this focus on one mode of action, the high proportion of refractory epilepsies has remained constant since the 1850s (*86, 87*). The underlying causes of these refractory epilepsies must be identified in order to develop more effective therapies. As the molecular basis of LD has been defined, we have now developed an AEF that targets its underlying cause. As molecular etiologies in other epilepsies become clear, precision therapies, possibly also utilizing the AEF platform, become feasible. LD is also a non-classical GSD. There are over 14 different types of GSDs characterized by glucan accumulations, and 1 in 20,000 people have some form of GSD though the individual types are considered rare (*7, 88*). While the glycogen in some GSDs is of normal structure like in Pompe disease, other GSDs such as Cori disease and Andersen disease are characterized by polyglucosans, like LD (*7*). Two AEF drugs (VAL-0417 and Fab-GAA) have now been shown to target, respectively, cytosolic polyglucosan (the present study) and normal glycogen in both the cytosol and lysosome (*47*). These drugs could be repurposed and/or modified to target glycogen in other GSDs. Not only is VAL-0417 the first drug with the potential for providing a significant clinical benefit to LD patients, it is an example of a precision therapy and expands the repertoire of antibody-based drugs that can be used to treat human disease.

## Materials and Methods

### Generation of the antibody-enzyme fusion VAL-0417

VAL-0417 was designed and produced by Valerion Therapeutics (Concord, MA) (Supplementary Methods). Activity of the purified protein was quantified by an amylase activity colorimetric kit (BioVision) utilizing the amylase-specific E-G7-pNP substrate (Fig. 1E (*51*).

### Mouse lines

*Epm2a-/- (8, 89)* and *Epm2b-/- (9)* mice have been previously described. C56Bl/6 WT, *Epm2a*-/- and *Epm2b*-/- animals were maintained in a 12:12 hr light-dark cycle and were given ad libitum access to food and water. All procedures were approved by the UK Institutional Animal Care and Use Committee (IACUC) as specified by the 1985 revision of the Animal Welfare Act.

### Purification of native LBs from *Epm2a*-/- and *Epm2b*-/- mice

19-24 month old *Epm2a*-/- mice were euthanized by CO2 and decapitation, and brain, heart and hindlimb skeletal muscle were immediately harvested, flash frozen and stored at −80°C. 12 month old *Epm2b*-/- mice were euthanized by cervical dislocation, and muscle tissues were similarly collected and stored. Tissues were pulverized over liquid N_2_ using a Freezer/Mill Cryogenic Grinder (SPEX SamplePrep). Powdered tissue was weighed and homogenized on ice in 4 vol. lysis buffer (100 mM Tris-HCl pH 8.0, 200 mM NaCl, 1mM CaCl2, 0.5% sodium azide) using a Dounce tissue grinder. For larger volumes, the grinding pestle was attached to a motorized drill. The homogenate was centrifuged for 10 min at 10,000×g at 4°C in a Ti-70 rotor and Optima XPN-90 ultracentrifuge (Beckman Coulter) and the supernatant was removed. The pellet was resuspended in an equivalent volume of lysis buffer, and 20% SDS was added for a final concentration of 0.2%, and 20 mg/ml Proteinase K (Invitrogen) was added for a final concentration of 0.4 mg/ml. Proteolytic digestion was performed in a 37°C water bath overnight. The digested samples were then syringe-filtered through 140 μm and 60 μm nylon net filters installed in Swinnex filter holders (Millipore), centrifuged at 16,000×g for 5 min, and the supernatant was removed. The LBs were resuspended in 10% SDS and then washed 5 times in LB buffer (10 mM HEPES-KOH pH 8.0, 0.1% sodium azide) each time with centrifugation at 16,000×g for 5 min. Final LB pellets were gently and thoroughly dispersed in LB buffer with a pipet. Polysaccharide yield at various steps of the purification were determined using the Pflüger method (Supplementary Methods). 2 μL of LB preparations were stained with 5 μL 20× Lugol’s iodine, mounted on glass slides with a glass coverslip, and visualized at 100x using a Zeiss Axioimager Z1 equipped with an AxioCam 1Cc5 color camera. Individual LBs in 3-5 micrographs were measured manually using ImageJ and a frequency distribution was calculated for LBs from each tissue type using Prism 6.0 software (GraphPad) and a bin width of 1 μm. Only clearly delineated LBs were measured; clumps and dust-like particles were excluded. Purified LBs were stored at −20°C. Iodine spectra and LB phosphate content of LBs were determined as described in Supplementary Methods.

### *In vitro* degradation assays

Corn starch used for degradation assays and microscopy was purchased from Sigma. Starch and LB degradation experiments were performed in degradation buffer (30 mM HEPES-KOH pH 7.5, 5 mM MgCl_2_, 5 mM CaCl_2_) and a total volume of 100 μL. Reactions were performed in triplicate using PCR strip tubes and moderate agitation on a vortex to keep substrates in suspension, and we found that this level of agitation did not affect enzyme activity. Degradation reactions were allowed to proceed overnight (13-18 hours) unless otherwise indicated, after which tubes were centrifuged to pellet undigested substrate. 50 μL of the supernatant (containing the soluble degradation product) was transferred to a new tube with 50 μL 2 M HCl and boiled for 2 hours at 95°C in a C1000 thermocycler (Bio-Rad) to hydrolyze all glucans to glucose. Samples were neutralized with 50 μL 2 M NaOH, and glucose was determined using the Boehringer-Manheim D-glucose kit (R-BioPharm). For visualization of LB degradation product, degradation reactions were allowed to proceed as above, and the reactions were centrifuged to pellet undigested substrate. After 75 μL of the supernatant was removed, the remaining 25 μL was resuspended, and 5 μL was stained with 5 μL 20× Lugol’s iodine, mounted on glass slides with a glass coverslip, and visualized using a Nikon Eclipse E600 using DIC/Nomarski contrast and an AxioCam MRm camera at 100x. In addition to the R Biopharm D-glucose kit, the absorbance-based PGO assay (Sigma) was also used to quantify glucose in starch degradation assays.

*In vitro* degradation of LBs in muscle homogenates was performed as follows: 300 mg of skeletal muscle from WT or laforin KO mice was pulverized over liquid N_2_ and homogenized in 4 vol. degradation buffer. Homogenate was split into 2 × 500 μL aliquots, and 25 μg VAL-0417 was spiked into one aliquot for a final concentration of 0.05 mg/ml. Samples were incubated on a rotator overnight at room temperature and the next morning centrifuged for 5 min at 16,000×g. The supernatant was removed, and the pelleted material containing LBs was resuspended in degradation buffer and boiled for 30 min on a 95°C heat block to solubilize LBs. The boiled samples were clarified by centrifugation (16,000×g for 5 min) and 25 μL of supernatant was added to a microplate. 50 μL of 1×Lugol’s iodine and 25 μL of water were added to samples, and absorbance was measured at 550 nm.

### Electron microscopy

For SEM, starch granules were incubated with enzymes in degradation buffer overnight while agitating, washed in 1mL 100% ethanol, dried in a vacuum centrifuge, and applied to carbon tape on a pin mount. Samples were coated with gold and platinum to improve sample conductivity, and visualized under high vacuum at 2 kV using an FEI Quanta 250 field emission scanning electron microscope. Washing in ethanol, drying, and manual application to carbon tape did not yield satisfactory results for LB visualization (fig. S4), so alternatively, diluted LBs were applied to a mounted silicon wafer, lyophilized, coated in gold and platinum, and visualized at 2 kV with the FEI Quanta 250.

### Profiling of VAL-0417 degradation product by HPAEC-PAD

80 μg of LBs were treated with 2.67 μg of VAL-0417 in degradation buffer to a final concentration of LBs at 1 μg/μL. The enzymatic reactions were done at 37°C in triplicates. 20 μL samples were removed at 24, 48, 72 hour intervals followed by addition of 2.67 μg of VAL-0417. The aliquots were stored at −20°C until they were run. The final reactions were continued at 37°C for total of 168 hours. All samples were profiled using CarboPac PA-100 column (Thermo-Dionex, 4 x 250 mm) and detected with PAD detector. 5 μL of samples from each time point (corresponding to 5 μg LBs total) were injected on the column. Blank LBs and degradation buffer were injected as controls. Glucose and maltose in the degradation reactions were quantified by comparing with standards of known amount (fig. S7A). 5 μg Maltrin100 was also included as a standard for oligosaccharide profiling (fig. S7E).

### Cell culture studies of VAL-0417 uptake and polysaccharide degradation

PTG and the catalytic alpha-subunit of protein phosphatase 1 (PP1Cα) were stably expressed in HEK293 cells to generate a stable cell line that accumulates polyglucosan (fig. S6 and Supplementary Methods). To determine the effect of VAL-0417 on polysaccharide degradation, cells were plated in 96-well plates at a density of 40,000 cells/well, and after 4 days various concentrations of VAL-0417 were added for 20 hr to fresh growth media. Cells were then washed 3 times with PBS, fixed for 7 min with 3.7% formaldehyde in 90% ethanol, washed again 3 times with PBS and polysaccharide content was determined by hydrolysis and glucose determination (Supplementary Methods). Statistical significance was assessed by using an unpaired Student t-test.

For Western analyses, 20 μg of total cell lysates were separated on SDS-PAGE and protein transferred to nitrocellulose membranes that were probed with antibodies against human pancreatic α-amylase (Abcam #ab21156) and ENT2 (Alomone Labs #ANT-052) followed by appropriate HRP-conjugated secondary antibodies and chemiluminescence.

### *In vivo* mouse studies

For each IM injection, 20 μL VAL-0417 (30 mg/ml; 0.6 mg per injection) or 20 μL PBS was injected into one gastrocnemius. At the indicated time points post-injection(s), mice were euthanized by cervical dislocation and decapitation. Tissues were harvested, flash frozen, powdered over liquid N_2_ using Freezer/Mill Cryogenic Grinder (SPEX SamplePrep), and stored at −80°C. Powdered tissues were weighed, homogenized in assay buffer from the BioVision amylase assay kit and protein concentration was determined using the Pierce BCA Protein Assay Kit (ThermoScientific). For the ELISA assay, tissue homogenates were diluted in PBS for a final concentration of 0.1 μg protein/μL in 100 μL. Diluted homogenates were added to 96-well plates for the ELISA utilizing a capture antibody binding the 3E10 Fab fragment and anti-AMY2A as the detection antibody (Supplementary Methods). Polysaccharide content was determined using the Pflüger method (Supplementary Methods).

ICV studies were performed by Northern Biomedical Research, Inc. (Spring Lake, MI). An ICV cannula attached to an osmotic pump via a catheter was surgically implanted in *Epm2a*-/- mice, and VAL-0417 (30 mg/ml) or PBS was continuously administered to the lateral ventricle for 28 days (0.11 μL/hr). On day 29, mice were euthanized by isofluorane/oxygen sedation and perfusion via the left cardiac ventricle with 0.001% sodium nitrite in heparinized saline. Brains were harvested, weighed, sectioned using a Rodent Brain Matrix (RBM-2000C, ASI Instruments) with 2.0 mm coronal section slice intervals, flash frozen and stored at −80°C. Slices were directly homogenized in BioVision assay buffer using polypropylene pellet pestles in microcentrifuge tubes. Protein and polysaccharide content were determined by BCA assay and the Pflüger method (Supplementary Methods), respectively.

Statistical significance of polysaccharide reduction was determined by one-way or two-way analysis of variation (ANOVA) using the Prism 6.0 software (GraphPad).

## Supplementary Materials

Materials and Methods

Fig. S1. Starch degradation with different VAL-0417:starch ratios.

Fig. S2. Iodine staining of LB purification fractions.

Fig. S3. Size distribution of LBs purified from different tissues and appearance of dust-like particles.

Fig. S4. Scanning electron micrographs of LBs from brain, heart and skeletal muscle, washed in ethanol, dried, and applied to carbon tape.

Fig. S5. Purification of LBs from skeletal muscle of 12 month-old *Epm2b*-/- mice.

Fig. S6. Polysaccharide levels relative to protein concentration in three cell lines: Rat1, HEK293, and HEK293 stably expressing PTG and PP1Cα.

Fig. S7. HPAEC-PAD chromatograms of control samples and additional time points from LB degradation with VAL-0417.

## Acknowledgments

We thank Drs. Craig Vander Kooi, Jeffrey Rush, and Ramon Sun of the University of Kentucky (UK) Department of Molecular and Cellular Biochemistry, Deb Ramsdell and Drs. Beth Goad, Rob Schaffer, and Hal Landy of Valerion Therapeutics for valuable feedback and discussions. We also thank Drs. Berge Minassian and Felix Nitschke for fruitful discussions and recommendations. We appreciate the technical support of Dr. Carole Moncman, Azin Akbari, Nico Briot, the UK Electron Microscopy Core, Dr. Thomas Wilkop and the UK Light Microscopy Core. We also thank Biswa Choudhury and Sulabha Argade for their assistance with HPAEC-PAD experiments (GlycoAnalytics, University of California, San Diego, CA).

## Author contributions

M.K.B., T.M., D.A., and M.S.G. conceived of the project and designed experiments. M.K.B., A.U., and G.A. performed experiments and analyzed data. G.A. designed and optimized the ELISA. D.M.S., A.D.P.R. and P.J.R. made stable cell lines, performed VAL-0417 uptake studies and analyzed data and provided *Epm2b*-/- tissues. J.J.M. performed intramuscular injections. B.L.H. and J.R.P. designed experiments and analyzed data. J.Z. oversaw ICV cannula implantations, injections and euthanasia. M.K.B. and M.S.G. wrote the paper.

## Data and materials availability

Researchers may obtain VAL-0417 with a material transfer agreement from Valerion Therapeutics. All reasonable requests for collaboration involving materials used in the research will be fulfilled provided that a written agreement is executed in advance between Valerion Therapeutics and the requester (and his or her affiliated institution). Such inquiries or requests should be directed to D.A.

