## Supplementary Material for "Targeting pathogenic Lafora bodies in Lafora disease using an antibody-enzyme fusion"

**Materials and Methods**

**Expression and purification of VAL-0417**

The cDNA encoding the human IgG1 Fab-linker-AMY2A heavy chain and light chain were synthetically produced with codon optimization for mammalian cell expression and cloned into pTT5. HEK293 cells expressing a truncated variant of the Epstein Barr Virus nuclear antigen 1 (HEK293-6E) increase the volumetric yield of monoclonal antibodies and fragments and were used for VAL-0417 expression (*90*). 2 1L cultures of HEK293-6E cells in 2L shake flasks were transfected with 1 mg of total plasmid DNA/L (1:1 ratio heavy chain:light chain) culture using PolyPlus linear Q-PEI at a 1:1.5 (w/v) DNA:PEI ratio. Culture parameters were monitored using a ViCell XR (Beckman Coulter) for density and viability. Culture was harvested 5 days post transfection via centrifugation for 5 minutes at 1000×g. The conditioned culture supernatant was clarified by centrifugation for 30 minutes at 9300×g.

Pre-packed CaptureSelect IgG-CH1 affinity columns (Fisher) were equilibrated in PBS (pH 7.2). VAL-0417 from 2 L of exhausted supernatant was top-loaded onto affinity columns (2 x 1 mL columns in tandem) at 4°C overnight. The column was washed with approximately 15 column volumes (CV) of PBS, 15 CV of buffer B (1×PBS with 500 mM NaCl, pH 7.2) and 15 CV of PBS. The resin-bound fusion protein was eluted with 10 CV of Buffer C (30 mM NaOAc, pH 3.5-3.6), collecting the protein in 1 mL fractions diluted in 1/10th volume Buffer D (3 M NaAcetate pH~9.0) to neutralize. To minimize the elution volume, elution was paused for several minutes between each fraction collected. Fractions were analyzed by A280 prior to pooling fractions and pools were analyzed by SDS-PAGE. VAL-0417 remained in the non-bound pool from the first affinity chromatography pass. The above procedure was repeated to capture remaining fusion protein. The affinity pools were combined prior to dialysis. The combined CaptureSelect IgG-CH1 affinity pool (18 mL) was dialyzed against 3 x 1 L of dialysis buffer (20 mM Histidine, 150 mM NaCl, pH 6.5) at 4°C. The dialyzed pool was concentrated to 1 mg/mL using a VivaSpin 20 (10K MWCO, PES membrane) centrifugal device prior to final analysis and storage at -80°C. Purified Fab-AMY was analyzed by size exclusion chromatography (Agilent HP1100) showing a single peak before and after a freeze-thaw cycle, indicative of a single, stable species.

**Pflüger method for polysaccharide isolation**

Glycogen and polyglucosan from *Epm2a*-/- and *Epm2b*-/- mice have been purified and characterized by multiple groups using the Pflüger method, but a chaotropic salt must be added to enhance the efficiency of polyglucosan precipitation (*20*). We wanted to clarify that under native conditions, and prior to the Pflüger treatment, LBs in *Epm2a*-/- and *Epm2b*-/- tissues are intact, micron-sized structures (Fig. 2 F to H, Fig. 3, and fig. S5B). Pflüger treatment converts the LBs to much smaller polysaccharide molecules that are on the same nanometer scale as glycogen molecules, but they contain elevated phosphate and an altered chain length distribution (*20, 21*). Thus, the term "LB" refers to the native, micron-sized polysaccharide-containing structures found in LD tissues, and "polyglucosan" refers to the abnormal polysaccharide comprising LBs, which is released with Pflüger treatment.

The Pflüger method is direct and very sensitive, and was used to track polysaccharide yield throughout the native LB purifications or to detect polysaccharide content of mouse tissues for *in vivo* studies. Aliquots from native LB purifications or tissue homogenates were added to 10 vol. 30% KOH, boiled for 2 hours, and allowed to cool. 2 vol. cold ethanol and 10 μL LiCl (1 M or 20 mM) were added, and samples were precipitated overnight at -20°C. Precipitated samples were centrifuged for 10 min at 16,000×g at 4°C, the supernatant was removed, and the polysaccharide pellet was resuspended in water. Two additional precipitations with cold ethanol and LiCl were carried out, each for 1-2 hours at -20°C. The final pellet was washed in cold ethanol and resuspended with vortexing in 200 μL water. Polysaccharide was quantified by overnight hydrolysis with amyloglucosidase from *Aspergillus niger* (Sigma) and glucose determination was carried out using the R Biopharm Inc. D-glucose kit (Fisher). Fluorescence (ex340/em445) rather than absorbance of the NADPH product (stoichiometric to glucose) was measured for higher sensitivity.

**Separation of glycogen and LBs in native LB purification**

After the initial centrifugation step in the LB purification, a fraction of the heart and muscle glycogen was consistently present in the supernatant (Fig. 2, B and C). In our preliminary purifications, we found that the homogenate and pellet fractions stained brown with Lugol's iodine after boiling, as did the final LBs and amylopectin, but the supernatants stained yellow like glycogen (fig. S2A). We also stained the Pflüger-purified polysaccharide fractions from our large-scale preparations in Figure 2C with Lugol's iodine, measured absorbance at 550 nm, and compared the values to the polysaccharide concentrations determined by hydrolysis and glucose measurement (fig. S2B). While the polysaccharide from the supernatant did not absorb at 550 nm, polysaccharide from the pellet, filtrate and final fractions produced a high absorbance relative to concentration. Polysaccharide in the homogenate fraction was less absorbent relative to concentration. These data indicate that the polysaccharide in the heart and muscle tissue homogenates included both glycogen and polyglucosan (in the form of LBs), that were separated into supernatant and pellet fractions after low-speed centrifugation. No polysaccharide was detected in the supernatant fractions from brain (Fig. 2, B and C). This result is consistent with the observation that in normal mice, glycogen levels are lower in brain than in other tissues in part due to its rapid catabolism after euthanasia (*91*).

**Iodine-based measurements**

Lugol's iodine was prepared as a 20×stock (1.5 M KI and 100 mM I_2_). To detect LBs in preliminary purifications, 50 μL samples from purification fractions were boiled for 15 minutes on a 95°C heat block, clarified by centrifugation (16,000×g for 1 min), and 35 μL of the supernatant was added to 50 μL 1×Lugol's iodine and 15 μL water for 100 μL total (fig. S2A). To analyze iodine absorbance of polysaccharide from LB purifications, Pflüger-isolated polyglucosan fractions were added to 50 μL 1×Lugol's iodine for 100 μL total and absorbance was measured at 550 nm. Absorbance was graphed alongside glucose-based concentrations to illustrate the fractionation of polyglucosan and glycogen into supernatant and pellet fractions, respectively (fig. S2B). For spectral scans, 50 μg LBs, rabbit liver glycogen (Sigma), and potato amylopectin (Sigma) were solubilized by boiling for 30 min, and added to 50 μL 1×Lugol's iodine for 100 μL total. Absorbance scans were performed at 400-800 nm in 10 nm steps.

**Determination of LB phosphate content**

Muscle glycogen and LBs contain covalent phosphate linked to the C2, C3, and C6 hydroxyls of glucose moieties, and the relative ratios of these modifications are equivalent (*21, 26, 92*). Boiling in mild HCl hydrolyzes glycosidic bonds and releases the acid-labile C2- and C3-linked phosphate leaving C6 phosphoesters intact in the form of glucose-6-phosphate (*21*). Thus, inorganic phosphate after mild acid hydrolysis represents only C2- and C3-linked phosphate, i.e. two-thirds of total phosphate. To determine phosphate content, skeletal muscle LBs were boiled for 2 hours at 95°C in 1 M HCl. Reactions were neutralized with NaOH and inorganic phosphate in the sample was determined using the Pi ColorLock Gold Phosphate Detection System (Innova Biosciences). Glycogen purified from rabbit skeletal muscle as previously described was used as a positive control, and the levels of phosphate detected using our method are consistent with two-thirds of published total phosphate levels (*26*).

**Generation of HEK293-PTG/PP1Cα cells and polysaccharide quantitation**

Protein Phosphatase 1 (PP1) stimulates glycogen synthesis by both dephosphorylating and activating glycogen synthase and inhibiting glycogen phosphorylase, the primary enzymes catalyzing glycogen synthesis and degradation, respectively (*93*). Protein targeting to glycogen (PTG) is a regulatory, glycogen-targeting subunit of PP1 that stimulates glycogen synthesis when overexpressed in cell lines (*94, 95*). Constitutive overexpression of PTG in WT mice has been shown to lead to cerebral polyglucosan accumulation that is similar to LBs in *Epm2b-/-* mice (*28*). We sought to design a cell line that accumulates polyglucosan like that which makes up LBs.

HEK293 cells (*96*) were co-transfected with plasmids pCDH-FLAG-PTG and pCDH-(HA)_3_-PP1Cα-GFP harboring mouse PTG and the human PP1Cα (catalytic alpha-subunit of PP1) cDNAs, respectively. Mixed clones were selected for ~10 days in the presence of 1 μg/ml hygromycin and 0.2 mg/ml puromycin, expanded and stored in liquid N2. Analyses of protein expression and glycogen synthase activity ratio in the absence and presence of glucose-6-phosphate indicated that both proteins were expressed and the glycogen synthase activity ratio was increased 3-fold, from 0.02 in control cells to 0.06 in transfected cells. Quantitation of the expression of PTG and PP1Cα is difficult because the basal levels are very low, undetectable under our conditions.

For quantitation of polysaccharide levels HEK293, HEK293-PTG/PP1Cα and Rat1fibroblasts were plated in 96-well plates at a density of 40,000 cells/well. After 48 hr, media was changed and the next day, the cells were washed 3 times with PBS followed by overnight incubation at 40°C in 50 μL of 0.2 M sodium acetate pH 4.8 containing 0.2% Triton and 0.3 mg/ml amyloglucosidase (*Aspergillus niger*; Sigma). Identical cell cultures were lysed in 50 μL of in 0.2 M sodium acetate pH 4.8 containing 0.2% Triton for protein determination by the Bradford procedure.

After overnight digestion, polysaccharide was measured by transferring 40 μL samples to a new 96-well plate. A 160 μL reaction mixture consisting of 0.375 M ethanolamine pH7.6, 5 mM MgCl_2_, 1.12 mM NADP, 2.5 mM ATP and 0.2 Units of G6PDH (Roche Biochemicals 10127655001) was added and OD340 nm was recorded. Subsequently, 0.75 units of hexokinase (Roche 11426362001) were added, the reaction incubated at room temperature for 30 min and OD340 recorded again. Background absorbance was subtracted from sample absorbance and glucose equivalents were determined based a digested glycogen standard curve. The engineered HEK293 line accumulated >30-fold more polysaccharide than the native HEK293 cells, and twice the level of normal glycogen found in Rat1 cells (fig. S6).

**Enzyme-linked immunosorbent assay (ELISA)**

The capture antibody raised against the 3E10 Fab fragment was generated by Valerion Therapeutics (Concord, MA). Wells of a 96-well plate were incubated with 100 μL of capture antibody overnight (~16 hours) at a final concentration of 2 μg/ml in PBS. All incubations were done in a humidified chamber. Wells were then rinsed 3 times with 200 μL PBS followed by incubation for 1 hour with 200 μL of blocking solution (5% non-fat milk in PBS). Wells were then rinsed 3 times with 200 μL PBS. 100 μL of diluted tissue homogenates were added and incubated for 1 hour. Wells were then rinsed 3 times with 200 μL of Tris-buffered saline (TBS) and 100 μL of primary antibody (anti-AMY2A, Abcam #ab21156) in 5% non-fat milk and TBS was added and incubated for 1 hour. The wells were then washed (solution added and incubated for 5 minutes before being removed) 3 times with TBS. 100 μL of secondary antibody (anti-rabbit IgG, HRP-linked, Cell Signaling Technology 7074) in 5% non-fat milk in TBS was added and incubated for 1 hour. Wells were then washed 5 times with TBS. 100 μL of TMB substrate (ThermoFisher N301) was added and a timer started. After the highest concentration of standard curve saturated or a pre-set time was met, the reaction was stopped by adding 100 μL of stopping solution (0.18% H_2_SO_4_). The plate was then read for absorbance at 450 nm.

**Supplementary Figures**

**Fig. S1.** Starch degradation with different VAL-0417:starch ratios. (A) Degradation with low ratio (10 μg VAL-0417, 1 mg starch). (B) Degradation with high ratio (10 μg VAL-0417, 100 μg starch).

**Fig. S2.** Iodine staining of LB purification fractions. (A) 50 μL samples were removed from homogenate, supernatant, and pellet fractions during preliminary LB purifications from skeletal muscle, boiled for 30 minutes, and clarified by centrifugation. 30 μL of the clarified sample was diluted with 15 μL water and stained with 50 μL 1x Lugol's solution in a microplate. Commercial liver glycogen and amylopectin were also stained as controls. (B) Concentration vs. iodine absorbance at 550 nm of Pflüger-isolated polysaccharide from skeletal muscle at each step of the purification scheme.

**Fig. S3.** Size distribution of LBs purified from different tissues and appearance of dust-like particles. Frequency distribution of LBs from each tissue type: (A) brain (total count: 473), (B) heart (total count: 247), and (C) skeletal muscle (total count: 252). Two fields of view from (D) brain and heart (E) LB preparations containing numerous dust-like particles are shown. Black boxes indicate areas that have been magnified and are labeled correspondingly (e1, e2, etc.). Occasionally very large bodies were observed (10-20 μm). They were difficult to distinguish from tightly packed clumps, so they were excluded from the size distribution histograms.

**Fig. S4.** Scanning electron micrographs of LBs from brain, heart and skeletal muscle, washed in ethanol, dried, and applied to carbon tape. Samples were visualized at 2kV under high vacuum by an FE Quanta 250.

**Fig. S5.** Purification of LBs from skeletal muscle of 12 month old *Epm2b-/-* mice. (A) LB purification yields. Mean ± SD of triplicate measurements are shown. (B) *Epm2b-/-* LBs stained with Lugol's solution and visualized by light microscopy. (C) Comparison of size distribution of skeletal muscle LBs from *Epm2a-/-* and *Epm2b-/-* mice. (D) Normalized iodine spectra of *Epm2b-/-* and *Epm2a-/-* skeletal muscle LBs. The spectral data are an average of 3 replicates.

**Fig. S6.** Polysaccharide levels relative to protein concentration in three cell lines: Rat1, HEK293, and HEK293 stably expressing PTG and PP1Cα. Data are expressed as the mean of 12 replicates ± SE.

**Fig. S7.** HPAEC-PAD chromatograms of control samples and additional time points from LB degradation with VAL-0417. (A) Overlay of control chromatograms: degradation buffer only (black line), LBs only (red line), and a mixture of 1 μg glucose and 1 μg maltose standards for quantitation (blue line). A small amount of maltose and low molecular weight oligosaccharides were detected in the LB control. Additional time points from the 168 hour LB degradation experiment with VAL-0417: (B) 24 hours, (C) 48 hours, and (D) 72 hours. Each chromatogram is representative of triplicate chromatograms. Small shifts in retention time (≤1 min) are typical between HPAEC-PAD runs. (E) Maltrin100, a mixture of low molecular weight oligosaccharides, was used as a standard for verifying degrees of polymerization (DP), i.e. number of glucose units.
