## Supplementary figures and images for "Targeting pathogenic Lafora bodies in Lafora disease using an antibody-enzyme fusion"

### Figure S1

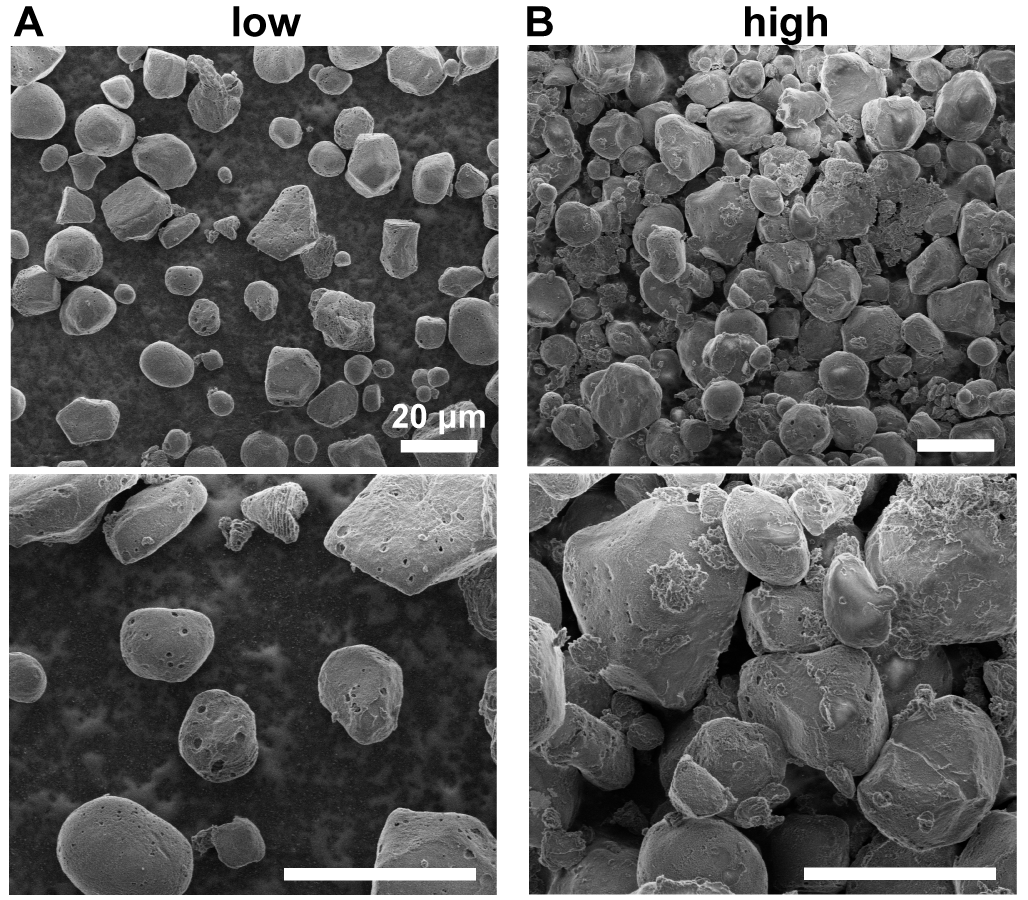

### Figure S2

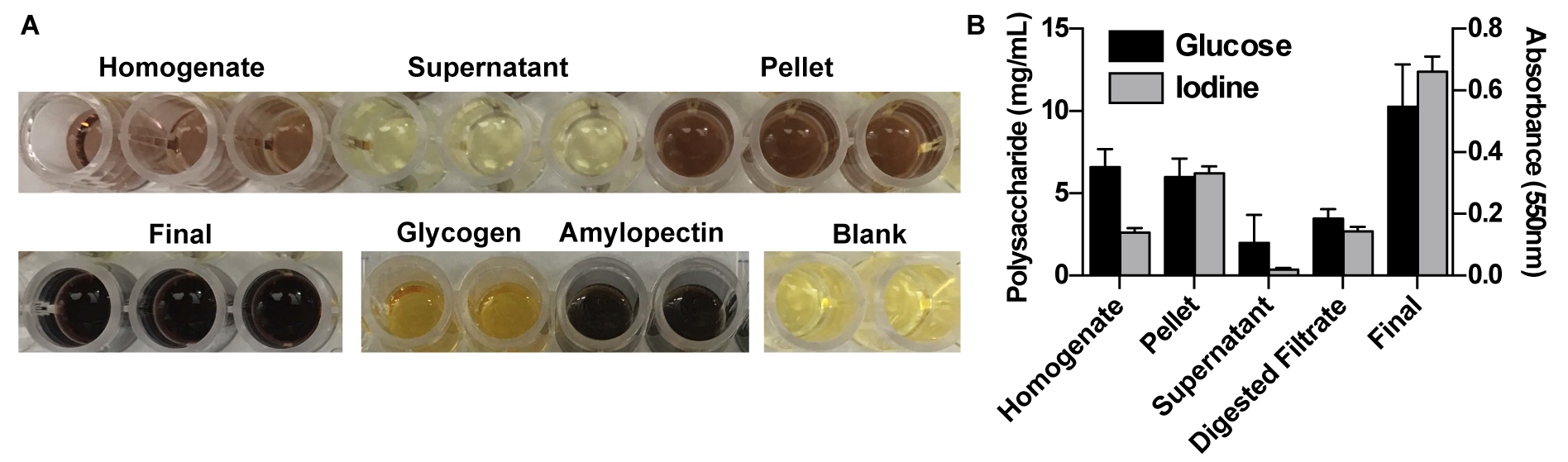

### Figure S3

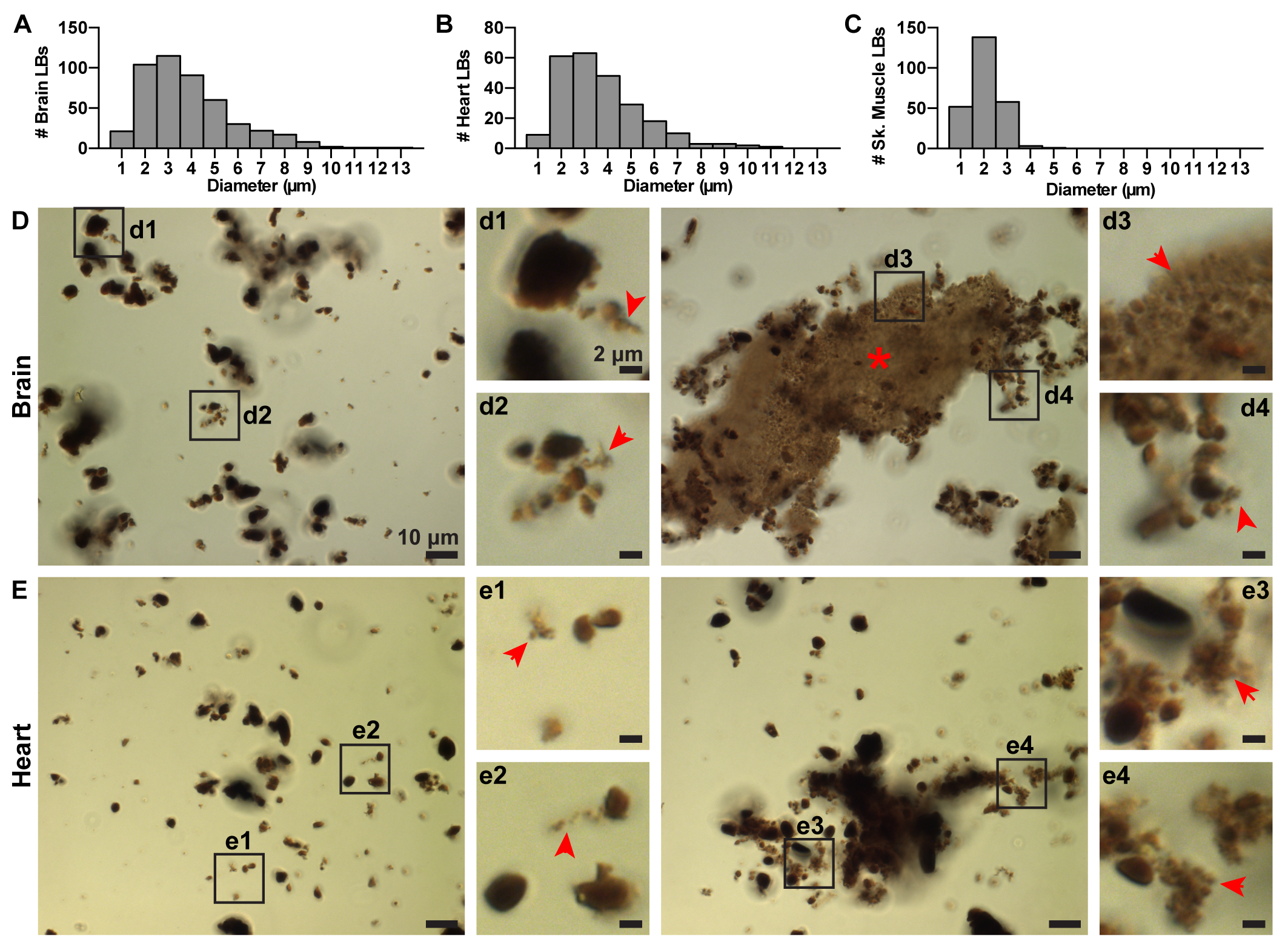

### Figure S4

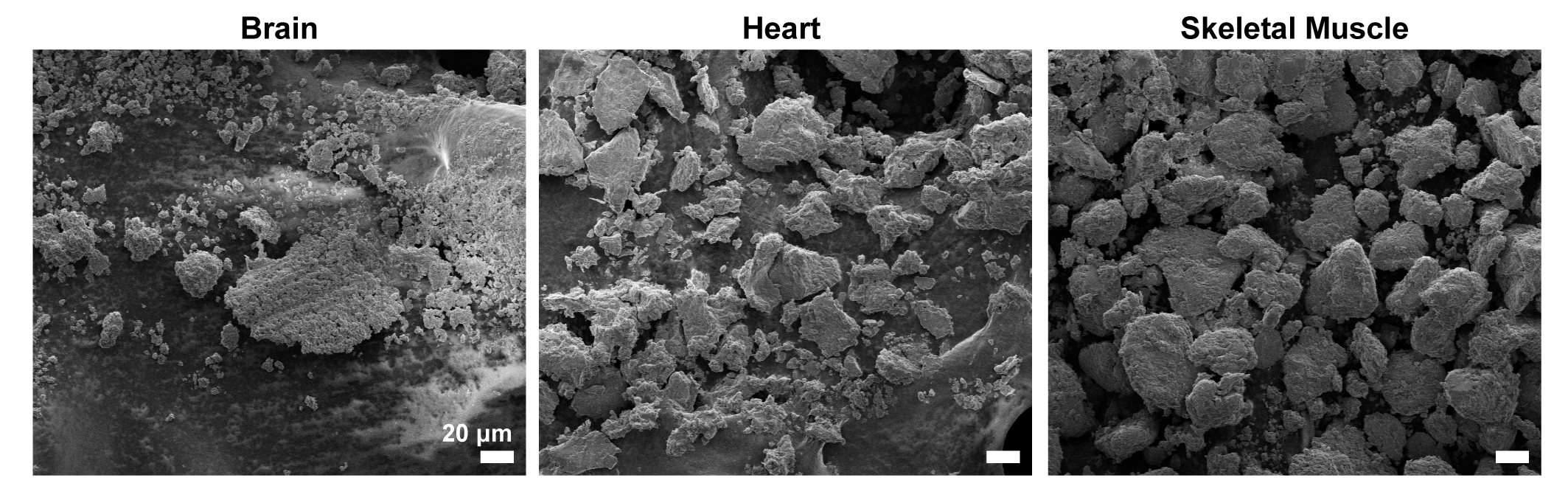

### Figure S5

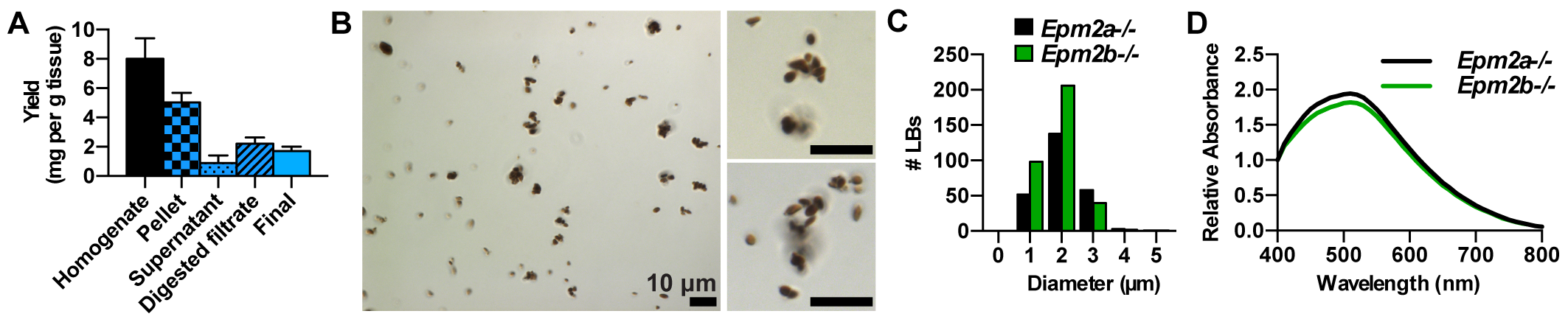

### Figure S6

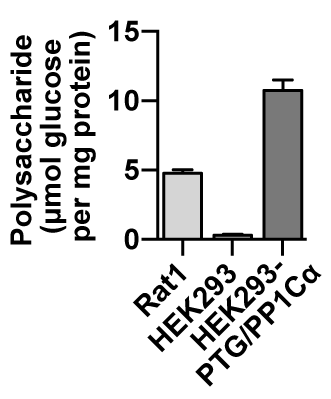

### Figure S7

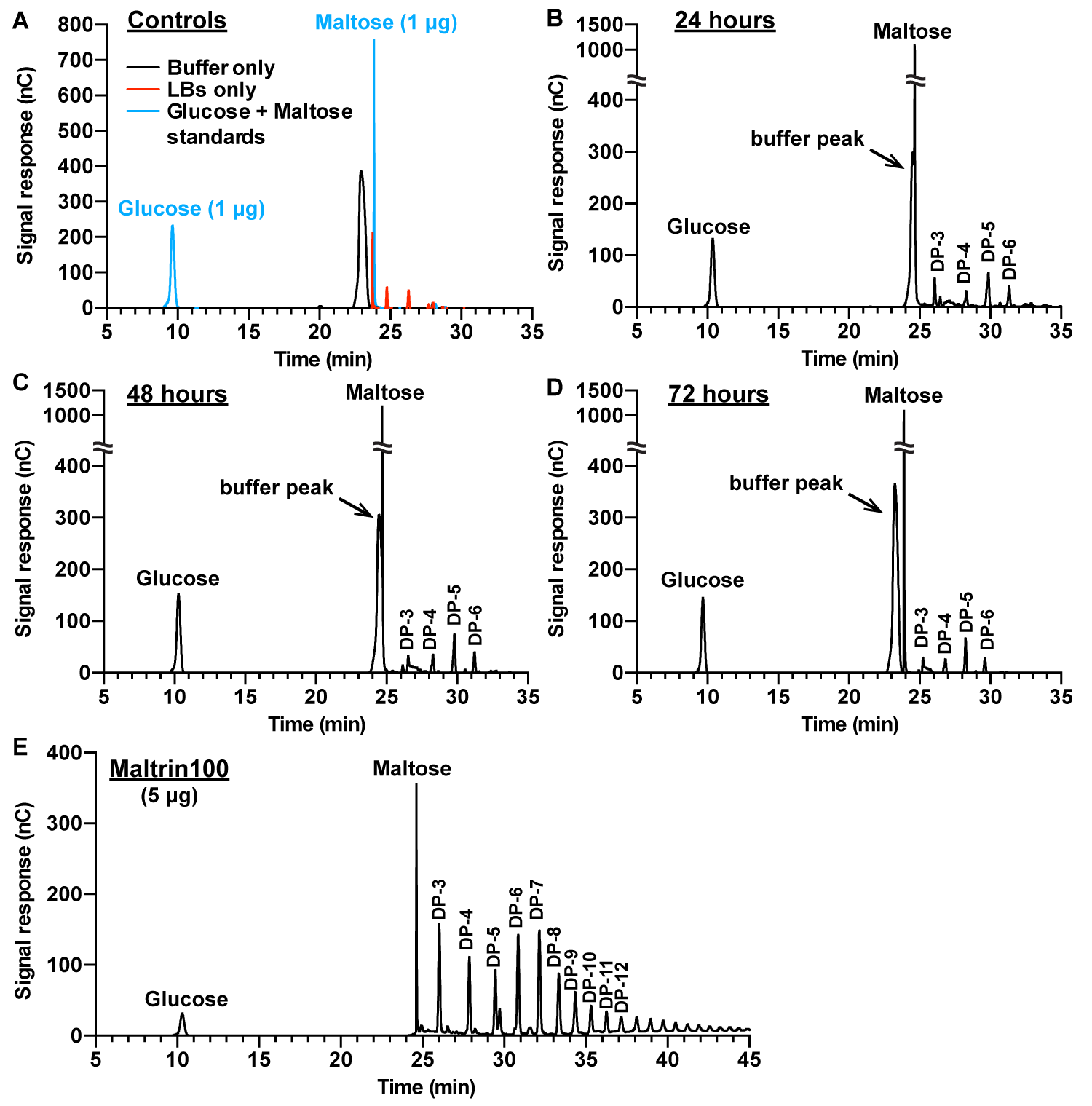
